# Genome-scale prediction of context-specific synthetic lethality beyond protein interaction networks

**DOI:** 10.64898/2026.07.31.742101

**Authors:** Pravin Baskar, Sahoo Parnika, Arnav Lakhdive, Saptarshi Bej, Sanu Shameer, Kamalakannan Vijayan

## Abstract

Identifying synthetic lethal (SL) interactions offers a principled framework for discovering disease-specific therapeutic targets. However, current machine learning approaches heavily rely on curated protein-protein interaction networks. Because these networks cover only ∼7,500 proteins, they severely restrict the search space of human gene pairs and introduce systematic biases toward well-characterized genes. To circumvent these limitations, we developed SLxGO, a network-independent machine learning framework that predicts SL interactions directly from semantic representations of Gene Ontology annotations encoded via BioBERT-derived embeddings. Benchmarked across multiple cross-validation schemes against eight state-of-the-art methods, SLxGO consistently achieved superior predictive ranking performance, maintaining robustness under cold-start conditions for previously unseen genes. Integrating cell line-specific transcriptional profiles extended this framework to context-dependent SL prediction across six distinct cell lines. Notably, we experimentally confirmed a context-specific *EFNA1*-*SLC29A1* SL interaction in HeLa cells, alongside synergistic pharmacological validation of an *ACVR1*-*SLC29A1* vulnerability. All predictions are hosted on SLiGO, an open-access database encompassing 30 million human gene pairs, establishing a comprehensive, genome-scale resource for context-specific vulnerability mapping across the human interactome.

## Introduction

Synthetic lethality (SL) describes a genetic interaction in which simultaneous disruption of two genes causes cell death, whereas perturbation of either gene alone is tolerated by the cell. This concept has emerged as a powerful framework for identifying cancer-specific vulnerabilities by exploiting genetic dependencies created by tumor-associated mutations. The clinical success of poly(ADP-ribose) polymerase (PARP) inhibitors in BRCA1/2-deficient cancers provided the first proof-of-concept for SL-based precision oncology and stimulated broad interest in systematically identifying SL interactions as actionable therapeutic targets (1–3)

Despite major advances in functional genomics, comprehensive experimental mapping of SL interactions remains impractical at genome scale (4). Although high-throughput approaches, including CRISPR, RNA interference, and chemical perturbation screens, can identify candidate SL pairs, the combinatorial space of human gene pairs exceeds 200 million, making exhaustive experimental interrogation infeasible (5). Moreover, SL interactions are highly context-dependent and can vary across tissue lineages, mutational backgrounds, and cellular states, such that interactions identified in one context may not be conserved in another, substantially increasing the experimental burden required to define clinically relevant SL landscapes (4, 6, 7).

Computational approaches have therefore become essential for prioritizing candidate SL interactions. Early methods, such as DAISY, inferred SL relationships using principles including mutual exclusivity of mutations, co-expression patterns, and functional similarity across cancer genomic datasets (8). Network-based approaches such as SLant (Synthetic Lethal analysis via Network Topology), the well-established random forest-based predictor behind the Slorth database, further incorporated conserved topological properties of protein-protein interaction (PPI) networks, including shortest path distances, shared neighbors, and centrality measures (9). More recently, machine learning approaches, including matrix factorization frameworks such as GRSMF (10) and SL2MF (11), and graph neural network approaches, such as GCATSL (12) and DDGCN (13), have improved predictive performance by integrating heterogeneous biological features and learning complex gene relationships from large-scale datasets. However, a major limitation persists across most existing ML-based approaches that depend heavily on curated PPI networks and knowledge graphs. Because these source networks lack consensus across databases (14) and skew heavily toward well-studied genes (5, 15), models are trained on highly biased data. For example, SLant can only predict SL pairs involving the ∼7,500 proteins present in the STRING PPI network it relies on, effectively excluding ∼13,000 human protein-coding genes. Such dependency restricts the scope of SL prediction and systematically limits the discovery of vulnerabilities across the genome. Beyond coverage limitations, many existing models also exhibit reduced generalizability when predicting context-specific SL interactions and interactions involving previously unseen genes, limiting their utility for identifying novel therapeutic vulnerabilities.

A recent comprehensive benchmarking study by *Feng et al.* (17) further highlights the limitation of many ML-based SL prediction methods. Across multiple datasets and evaluation strategies, all existing methods were reported to perform poorly based on ranking-based metrics. Classification performance of models were also observed to decline significantly under stringent evaluation strategies where predictions were made on unseen genes. This reduced generalizability stems from the inability of existing approaches to infer relationships for genes with little or no prior interaction information, thereby limiting their applicability to poorly characterized genes.

To address these limitations, we developed SLxGO, a network-independent machine learning framework that predicts SL interactions directly from semantic representations of Gene Ontology (GO) annotations trained on partially curated interactomes. SLxGO uses BioBERT-derived embeddings (16) to derive functional relationships among genes and infers network-like topological features without requiring experimentally derived interaction maps. Following the comprehensive benchmarking framework of *Feng et al.* (17), we evaluated SLxGO against eight state-of-the-art SL prediction methods across multiple cross-validation settings and negative sampling strategies. SLxGO consistently outperformed existing approaches, including in cold-start scenarios involving previously unseen genes. To capture context-dependent biology, we integrated cell line-specific gene expression profiles to model SL interactions across six cell lines, resulting in the first prediction and experimental validation of two novel SL pairs: *EFNA1*-*SLC29A1* and *ACVR1-SLC29A1*. Finally, to facilitate community access, we developed SLiGO, an open-access database containing genome-scale SL predictions across 30 million human gene pairs, substantially expanding the landscape of candidate disease vulnerabilities beyond existing interaction maps.

## Materials and Methods

### GO annotation and generation of BioBERT embeddings

To capture a comprehensive description of every human gene, the Wikigenes gene symbol along with GO annotations related to their PANTHER protein class, PANTHER protein family, molecular function, biological process, and cellular localization were collected from the Gene Ontology Resource Database (18, 19). These six annotations were fed into BioBERT, a domain-specific adaptation of the BERT language model pre-trained on a massive biomedical corpora (16), to generate embeddings for each gene. BioBERT was implemented using the **`NLU v 5.4.1`** released by John Snow Labs. By concatenating the six embeddings, a single 4,608-dimension vector was generated for each gene. A pair of 4,608-dimension vector representations, henceforth to be referred to as X_GO, was generated for each gene pair. A Python code for the pipeline is available at https://github.com/sshameer/SLxGO2026.

### Generating SLant network features and embeddings

The SLant algorithm and the associated PPI network were reproduced using R scripts available in the publicly accessible SLant Bitbucket repository (https://bitbucket.org/bioinformatics_lab_sussex/slant/src/master/). Minor modifications were made to the code to address incompatibility issues between the **`dplyr v 1.1.4`** package and the **`R 3.4`** environment. The SLant algorithm was used to generate 49 network features (such as centrality measures, number of n-hop neighbours, etc) for all gene pair combinations available within the SLant PPI network i.e. 9,247,150 gene pairs. Of these, 179,739 gene pair entries for which one or both genes could not be mapped onto wiki gene IDs were filtered out. Finally, a list of 9,067,411 gene pairs along with a 49-dimension vector representing their SLant network features (hence-forth to be referred to as X_SLant) mapped to the respective WikiGenes IDs were saved as a CSV file. An R script for this process described is available at https://github.com/sshameer/SLxGO2026.

### Network feature approximation using an artificial neural network

The SLant PPI network contains only 4,301 protein-coding genes. In order to predict X_SLant equivalent network features for gene pairs where one or both genes are absent from the PPI network, an artificial neural network (ANN) trained on the 9,216 dimension X_GO as input and min-max normalized 49 dimension X_SLant as output was developed using the **`PyTorch v2.5.1`**. Predicted normalized X_Slant features are finally rescaled to its original values. The architecture included two densely connected hidden layers of 2048 and 1024 neurons. Each layer used a Rectified Linear Unit (ReLU) activation function to introduce non-linearity, allowing the network to model complex patterns while maintaining computational efficiency. Mean imputation for data filling using the imputer function available in the **`Scikit-learn v 1.7.2`** package was used to ensure a complete input space for model training. A standard 80:20 train-test split was employed for training the ANN. The model was trained for 10 epochs at a learning rate of 0.0001, with each run randomising the order of input pairs. Training minimized for the mean squared error (MSE) loss function to learn the transformation. An IPython notebook describing the entire process is available at https://github.com/sshameer/SLxGO2026.

### Feature integration and Gradient Boosting classification

Principal Component Analysis (PCA) was performed on embeddings of five GO annotation types (excluding the Wikigenes ID encodings) and ten principal components were retained from each feature, resulting in a development of a 50-dimensional vector representation for every gene (henceforth to be referred to as X_GO_PCA). For each gene pair, the 50-dimensional representations of both genes were concatenated following both the alphabetic order of their gene names (referred to as ‘forward’ run) and the reverse order, yielding two 100-dimensional vector representations of GO features. Both variants of the 100 dimensional vector gene pair representations were further concatenated with the respective X_SLant representations to create vectors representing both gene annotation and gene network representation for any gene pair. The resulting 149-dimension embeddings were used as input for a Gradient Boosting classifier (a high-performing ensemble-based classifier) to predict whether a specific gene pair shows an SL interaction or not. The algorithm was trained using both variants of a gene pair representation, and their average score was presented as a cumulative confidence. The gradient boosting algorithm was executed using the **`HistGradientBoostingClassifier`** function available in **`Scikit-learn v 1.7.2`**, a faster version of classical gradient boosting, which does not compromise the quality of the model. Parameters were optimised for F1 score using the **`RandomizedSearchCV`** function, also available in the **`Scikit-learn v 1.7.2`**. IPython notebooks for the processes described are available at https://github.com/sshameer/SLxGO2026.

### Training and test dataset construction

Training and evaluation dataset required for SLxGO and other SL predictors was gathered from a previous-published benchmarking study evaluating ML-based SL predictors (17). This dataset consisted of 26,220 experimentally validated SL positive gene pairs sourced from SynlethDB database and 106,280 SL negative data estimated from DepMap transcriptomics data (20). In order to balance the positive and negative data during benchmarking runs, random sampling was performed on the latter and only 26,220 SL negative pairs were selected. The release version of SLxGO was trained on this balanced dataset (Dataset 1).

### Comparing the performance of SLxGO against SLant

Of the 52,440 (positive and negative) SL gene pairs in the balanced dataset, only 15,230 pairs could be found within the SLant PPI network, of which 8,885 were positive and 6,345 were negative SL gene pairs. In order to evaluate the performance of SLxGO directly with SLant, a new balanced set of the 12,690 SLant-compatible gene pairs (Dataset 2) was prepared for comparison of predictive capabilities between SLxGO and SLant.

### Benchmarking and Evaluation Strategy

In order to comprehensively evaluate the performance of our model, the *Feng et al.* (17) benchmarking study comparing eight SL prediction tools; namely GRSMF (10), SL2MF (11), GCATSL (12), DDGCN (13), SLMGAE (21), NSF4SL (22), KG4SL (23), and PTGNN (24); was extended to include the SLxGO algorithm. Datasets for benchmarking were gathered from the previous study (17). SL positive gene pairs in this dataset consisted of experimentally identified SL positive gene pairs published on SynLethDB (25). As previously described (17), three different strategies were used to generate the SL negative gene pairs found in the training and test data.

1. In ‘dependency-based sampling’ strategy, negative gene pairs were selected using cell line-specific gene dependency profiles, ensuring non-essential or non-synergistic gene combinations.
2. In ‘expression-based sampling’ strategy, genes with dissimilar expression profiles were assumed to lack functional interaction and were used to form negative pairs.
3. In ‘random sampling’ strategy, random gene pairs were selected from the gene pool while ensuring they did not overlap with known SL interactions.

*Feng et al.* (*17*) also relied on three different training and test data splitting strategies (CV_1_, CV_2_, and CV_3_) to perform a robust benchmarking, which was replicated in this study as well.

1. The CV_1_ (SL pair-level split) strategy involved randomly splitting the positive and negative SL data into training and test data such that both genes present in a test set gene-pair can be found in the training data gene-pairs.
2. In the CV_2_ (single gene-level split) strategy, training and test data were split such that only one of the genes in the gene-pair set in test data is present in the training set gene-pairs.
3. In CV_3_ (double gene-level split), training and test data were split such that neither of the two genes in test data gene-pairs could be found in training data. This simulates prediction scenarios for newly studied or poorly annotated genes.

To maintain consistency in benchmarking, model performances were evaluated using both classification and ranking metrics as proposed by *Feng et al.* (*17*). Standard classification metrics: Area Under the Receiver Operating Characteristic Curve (AUCROC), Area Under the Precision Recall Curve (AUPR) and F1 score, were used to evaluate the ability of the ML algorithms to correctly classify gene pairs as SL or non-SL. Similarly, NDCG@10, Precision@10 and Recall@10 metrics were used to evaluate the ability of the models to correctly rank SL predictions. All calculations were made using the metrics submodule of python **`Scikit-learn v 1.7.2`** package. For each training and test data split strategy, a classification score *C_datasplit_* was calculated by finding the mean of all three classification metrics (AUCROC, AUPR and F1 score). A classification score was assigned to each model following equation provided,

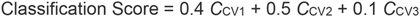

Similarly, for each training and test data split strategy, a ranking score R was calculated by finding the mean of all three ranking metrics (NDCG@10, Precision@10 and Recall@10), and a ranking score was assigned to each model based on the following equation,

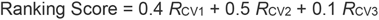

The rationale given behind a slightly higher weightage for CV_2_ was attributed to it representing a more realistic scenario (17). The same logic was applied for both ranking and classification scores. Finally, an overall score was also calculated for each model from the average of both classification and ranking scores.

### Cell line specific embeddings

To incorporate cell line specific features, gene expression data of 1,206 human cell lines were collected from The Human Protein Atlas repository (26). K-means clustering of the cell lines was performed using the **`Scikit-learn v 1.7.2`** package to identify genes that may not be globally important but are specifically important within particular cell lines. An elbow break method implemented using the **`kneed v 0.8.5`** package was applied to identify the best cluster size of 8 for K-means clustering. Principal Component Analysis (PCA) was performed using the **`Scikit-learn v 1.7.2`** package to identify principal components that represent 95% of the variance in gene expression among these clusters. The resulting 3524-dimension feature space was further reduced using Uniform Manifold Approximation and Projection (UMAP) dimensionality reduction using **`umap-learn v 0.5.3`**to generate pseudo genes representations of each cell line. The final representative pseudo gene set was selected using a trial and error method based on metrics such as trustworthiness (T), k-nearest neighbours (kNN) preservation (K), distance correlation (C), and variance ratio (V). The final embedding size of 16 was chosen using the formula below.

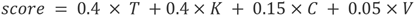

All metrics and the final score calculations were made by custom functions that are available at https://github.com/sshameer/SLxGO2026. Cell line specific embeddings of all 1,206 cell lines were generated (X_CL) and saved for use during run-time.

### Cell line-specific model SLxGO+

Limited by the size of cell-line specific labelled data, we considered only 6 cell lines with at least 100 positive (SL) pairs and 100 negative (NSL) pairs known fromSynLethDB (Dataset 3). The distribution of SL and NSL pairs in Dataset 3 is provided in Table 1.

**Table 1:** Distribution of reported synthetic lethal (SL) and non-synthetic lethal (NSL) gene pairs across six human cell lines].

| Cell line | SL | NSL |
| --- | --- | --- |
| HEK293 | 326 | 2,148 |
| A375 | 1,086 | 1,411 |
| A549 | 1,275 | 8,497 |
| HeLa | 1,254 | 4,804 |
| Jurkat | 2,595 | 64,966 |
| K562 | 8,655 | 83,637 |

For the six cell lines depicted in **Table 1**, gene information (X_GO) along with Network information (X_SLant) were concatenated with cell line specific embeddings (X_CL), resulting in the generation of a pair of 165-dimensional cell-line specific embeddings for a gene pair. Similar to the classic SLxGO version, the cell line specific version (SLxGO+) is also trained using both variants of the input vector, and uses **`HistGradientBoostingClassifier,`** a gradient boosting classifier from **`Scikit-learn v 1.7.2`**. Parameters were optimised for precision score using the **`RandomizedSearchCV`** function. The scores from using forward and reverse variants of the input vector are averaged to present the cumulative confidence score. The code for this implementation is available at https://github.com/sshameer/SLxGO2026.

## Database

In order to enable the discovery of SL and analysis of SLxGO and SLxGO+ results, a web-based user-friendly platform named SLiGO was established. The website is a lightweight full-stack implementation, with its frontend built using **`HTML5`**, **`CSS3`**, and **`JavaScript`**, while the backend uses Python packages such as **`Flask v 2.2.2`** for web framework and API handling, and **`Duckdb v 1.5.2`** as the primary database engine for storing the SL interaction data as a SQL lookup file.

The frontend communicates with Flask API endpoints and the backend queries the SQLite database to return JSON responses, cached results, and indexed searches, enabling efficient handling of the data to provide a fast, scalable, and user-friendly interface. The two key API are Gene Search and Gene Pair Search, which help in querying the desired results based on the given confidence interval.

The database supports searching, browsing and downloading SL interactions discovered by the SLxGO and SLxGO+ pipelines. SLiGO is open to the general public and can be accessed via https://databases.iisertvm.ac.in/sligo/ without needing to register or log in.

### Pathway activity scoring and relative dependency analysis for ephrin signaling, nucleoside salvage pathway and *de novo* nucleotide synthesis

Gene expression data for HeLa, HEK293, A549 and Jurkat cell lines were obtained from the Human Protein Atlas dataset (27). Expression values were log-transformed and normalized prior to analysis. To enable comparison across cell lines, gene-wise Z-score normalization was performed across the selected cell lines. For each gene *g* in cell line *i*, the *Z*-score was calculated as:

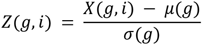

where *XX*(*g*, *i*) represents the normalized expression value of gene g in cell line i, and *μ*(*g*) and *σ*(*g*) denote the mean and standard deviation of gene g across the three cell lines.

Pathway activity scores for ephrin signaling, nucleoside salvage and *de novo* nucleotide synthesis were computed by averaging the *Z*-scores of genes associated with each pathway. Pathway-associated gene sets were downloaded from the Molecular Signatures Database (MSigDB) (28), including Reactome (CP:REACTOME) and KEGG (CP:KEGG) pathway collections. Among these, genes expressed in each of the four cell lines were identified, and only genes commonly expressed across all four cell lines were retained for subsequent analyses. The pathway-specific genes retained after expression filtering across all four cell lines are listed below.

Ephrin signaling: *EFNA1*, *EPHA2*, *EPHA4*, *EPHB2*

Nucleoside salvage pathway: *SLC29A1*, *SLC29A2*, *DCK*, *TK1*, *NT5C2*

*de novo* nucleotide synthesis: *CAD*, *DHODH*, *UMPS*, *RRM1*, *RRM2*

For each pathway *p* in cell line *i*, the pathway activity score was calculated as:

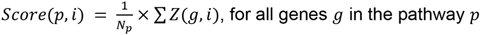

where *N_p_* represents the number of genes in the pathway *p*.

To quantify relative pathway dependence, a pathway dependency score was defined as the difference between nucleoside salvage and *de novo* synthesis pathway activities:

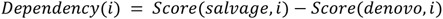

Positive values indicate greater reliance on nucleoside salvage pathways, whereas negative values indicate dominance of de-novo nucleotide synthesis.

### siRNA-mediated knockdown and chemical perturbation assays

HeLa and A549 cells were maintained in Dulbecco’s Modified Eagle Medium (DMEM; Gibco, 10569-010) supplemented with 10% fetal bovine serum (FBS) and 5% penicillin-streptomycin at 37 °C in a humidified incubator with 5% CO₂. Cells were passaged upon reaching 80-90% confluency.

For knockdown experiments, cells were seeded in 24-well plates to achieve ∼50% confluency at the time of transfection. Gene-specific siRNA duplexes targeting EFNA1 (Eurogentec, Belgium) and a non-targeting scrambled siRNA control were transfected using Lipofectamine RNAiMAX (Invitrogen, 13778150) following the manufacturer’s protocol. Briefly, siRNA (final concentration 1 µM) and RNAiMAX were separately diluted in Opti-MEM, incubated for 5 min, combined, and allowed to form complexes for 15 min prior to addition to cells in antibiotic-free medium. After 6 h, the medium was replaced with a complete growth medium, and cells were incubated for 24-48 h.

Knockdown efficiency was assessed by flow cytometry. At 48 h post-transfection, cells were fixed in 4% paraformaldehyde, permeabilized with 0.1% Triton X-100, and blocked with 5% bovine serum albumin. Cells were incubated with an anti-EFNA1 primary antibody (Invitrogen, 34-3300), followed by an Alexa Fluor-conjugated secondary antibody (Invitrogen, A21-441). Samples were analyzed using a BD FACS Lyric system.

For functional assays, cells were treated 24 h post-transfection with the ENT1 inhibitor Nitrobenzylthioinosine (NBMPR, HY-W010936) at 10 µM or 20 µM, or Dimethyl sulfoxide (DMSO) control (Sigma-Aldrich). Drug-containing medium was refreshed every 24 h for a total treatment duration of 72 h.

Cell viability was assessed using a crystal violet assay. Cells were fixed with 4% paraformaldehyde, stained with 0.5% crystal violet, washed, and air-dried. The images were obtained using a Nikon DSLR camera.

### Combinatorial chemical perturbation and synergy analysis

For combinatorial pharmacological validation of the predicted *ACVR1*-*SLC29A1* interaction, single-agent dose-response profiles were first established for the ENT1 inhibitor NBMPR and the *ACVR1* inhibitor ALK2-IN-2. Cells were seeded in 96-well plates and subjected to a fixed-dose combination assay, receiving a constant concentration of NBMPR (5 µM) alongside an escalating dose gradient of ALK2-IN-2 (0.125 µM to 20 µM). Equivalent volumes of DMSO were utilized across all wells to normalize vehicle concentrations. Cell viability was assessed via MTT assay at 72 h post-treatment.

### Quantification of Synergy via the Bliss Independence Model

To determine whether the pharmacological interaction between ALK2-IN-2 and the ENT1 inhibitor (NBMPR) was synergistic, additive, or antagonistic, cell viability datasets were evaluated using the Bliss Independence model (29). The fractional inhibition (fraction affected, E) for each single-agent treatment and their combination was calculated relative to the untreated vehicle control as described:

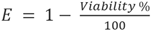

The theoretical additive effect expected from an independent, non-interacting relationship between the two compounds was computed:

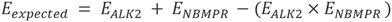

where *E_ALK_*_2_ represents the fraction affected by a given concentration of ALK2-IN-2 alone, and *E_NBMPR_* represents the fraction affected by the fixed concentration of 5 µM NBMPR alone. For the 0.125 µM baseline evaluation where single-agent titration data was absent, *E_ALK_*_2_ was defined as 0.

The net Bliss Synergy Score was calculated as the excess observed effect over the theoretical expectation:

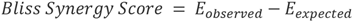

A Bliss Synergy Score greater than zero indicates a synergistic interaction, with a threshold of > 0.10 defined as biologically significant synergy.

## Results

### Overview of the SLxGO framework

To overcome limitations of ML-based SL prediction algorithms, i.e., their inherent dependence on curated protein-protein interaction (PPI) networks, suboptimal ranking performance and lack of interpretability, we developed SLxGO, a machine learning framework for genome-scale prediction of synthetic lethality directly from GO semantics (**Figure 1**). The architecture employs a three-stage pipeline: (i) generation of 4,608-dimensional functional embeddings for each gene using BioBERT; (ii) a neural network-based mapping of these semantic features to 49 topological features derived from the SLant framework (9); and (iii) integration of these features into a gradient-boosted classifier for final SL prediction. The second stage of the architecture effectively reconstructs proxy interaction networks from functional annotations, enabling genome-scale interrogation independent of experimentally determined interaction maps.

**Figure 1.**
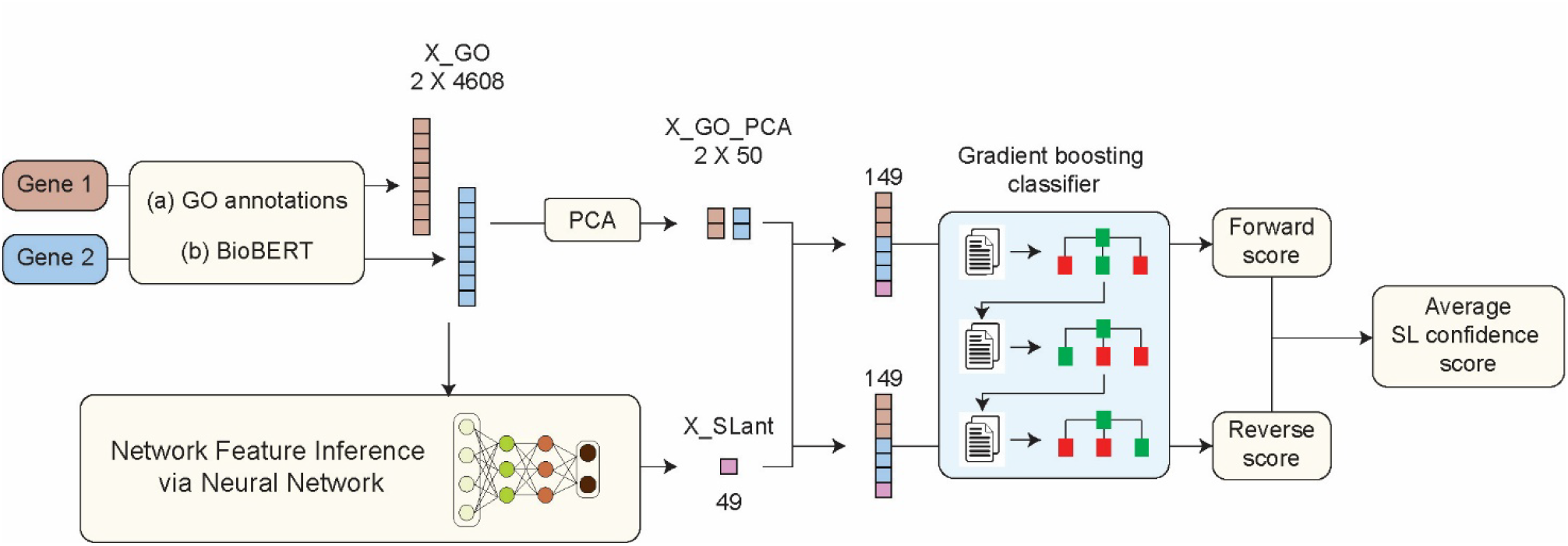
Architecture of the SLxGO framework. GO annotations of gene pairs were encoded using BioBERT to generate semantic gene embeddings. A neural network was trained to infer SLant-derived topological features from these embeddings, enabling reconstruction of proxy interaction networks independent of curated PPI maps. Inferred network features were combined with PCA-reduced GO embeddings and used as input to a gradient-boosted classifier for prediction of synthetic lethal interactions. Predictions are performed in both forward and reverse gene orientations, and the average of the two scores is reported as the final synthetic lethality confidence score.

### SLxGO Outperforms SLant in Predictive Performance

We next compared SLxGO with the SLant (9) framework using a balanced dataset of 12,690 gene pairs restricted to the SLant interaction network (Dataset 2 described in methods). SLxGO outperformed SLant across all classification metrics, achieving an AUCROC of 0.9784 compared with 0.9315 for SLant, together with improved accuracy (93.38% versus 86.12%) and F1 score (0.9339 versus 0.8599) (**Table 2, Figure S1**). These results indicate that integrating GO-derived semantic embeddings with inferred topological features provides a stronger predictive signal for synthetic lethality than conventional network-derived representations.

**Table 2.** Performance comparison of SLant and SLxGO on PPI network-covered gene pairs.

| Tool | Precision | Recall | F1 score | Accuracy | AUCROC |
| --- | --- | --- | --- | --- | --- |
| SLant | 0.868 | 0.852 | 0.860 | 0.861 | 0.932 |
| SLxGO | 0.941 | 0.939 | 0.940 | 0.940 | 0.984 |

To test the utility of SLxGO for poorly characterized genes, we evaluated the model on 39,750 SL gene pairs from Dataset 1 that lacked representation within the SLant PPI network and are therefore inaccessible to the network-dependent method. On this pseudo-validation set, SLxGO maintained strong predictive performance, achieving a precision of 0.8980 and an AUCROC of 0.9038 (**Table 3**). Although recall was moderately reduced 0.7372 (to prioritise precision), the overall model stability (F1 = 0.8097) demonstrates the capacity of SLxGO to extend SL prediction into the underrepresented genes of the genome, for which experimental interaction data remain sparse or absent.

**Table 3.**
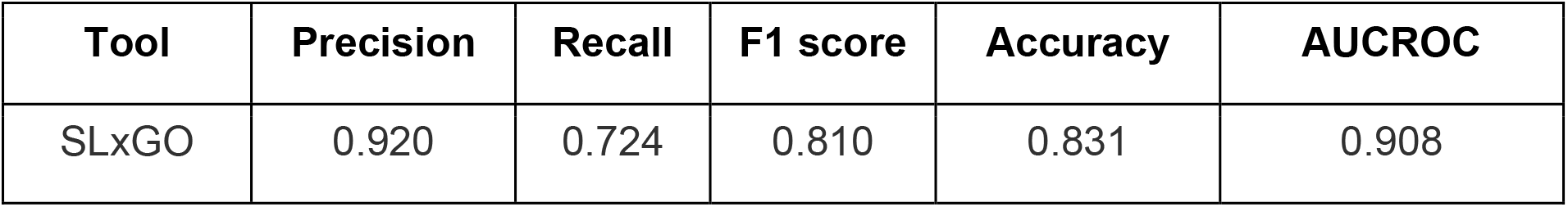
Performance of SLxGO on SL-positive gene pairs absent from the SLant interaction network.

| Tool | Precision | Recall | F1 score | Accuracy | AUCROC |
| --- | --- | --- | --- | --- | --- |
| SLxGO | 0.920 | 0.724 | 0.810 | 0.831 | 0.908 |

### Feature contribution analysis reveals the drivers of SL prediction

To interpret the features driving SLxGO predictions, we performed SHAP (Shapley additive explanations) analysis on the trained model. The 149 input features were grouped into three categories: network-derived features (49 dimensions), Gene1 GO features (50 dimensions), and Gene2 GO features (50 dimensions). In the training set, network-derived features contributed 12.7% of total model signal, while Gene1 and Gene2 GO features accounted for 42.6% and 44.6%, respectively (**Figure S2A and S2B**). Due to the sheer size of the test data, bootstrap sampling for 200,000 gene pairs across 10 iterations were performed and SHAP analyses were completed on each sample. Comparable contributions were also observed in the test case (13.4%, 43.7%, and 42.9% for network, Gene1, and Gene2 features, respectively) (**Figure S2C and S2D**), confirming the consistency and stability of feature contributions across partitions. The similar contributions of Gene1 and Gene2 features indicate that the model utilizes functional information from both genes in a pair without directional bias. Further analysis across major GO annotation categories as biological process, molecular function, cellular localization, and PANTHER protein class and family confirmed that GO-derived semantic features from both gene partners collectively dominate the predictive signal.

### Benchmarking SLxGO against state-of-the-art ML-based SL prediction models

To comprehensively evaluate the performance of SLxGO, we benchmarked our model against eight state-of-the-art ML-based SL prediction methods: GRSMF (10), SL2MF (11), GCATSL (12), DDGCN (13), SLMGAE (30), NSF4SL (22), KG4SL (23), and PTGNN (24).

The benchmarking framework followed the evaluation protocol described by *Feng et al* (*17*)., using identical datasets, data partitions, negative sampling strategies, and evaluation metrics. Benchmarking was conducted under three cross-validation schemes: CV_1_ (SL pair-level split, where both the gene per test pair is present in training), CV_2_ (single gene-level split, where one gene per test pair is absent from training), and CV_3_ (double gene-level cold-start split, where neither gene in a test pair appears in training). Each scheme was further evaluated using three negative sampling strategies: expression-based, dependency-based, and random. We successfully reproduced the previously published performance of all eight benchmark models. (**Supplementary Data 1**). Across both CV_1_ and CV_2_ settings, SLxGO consistently achieved the highest classification performance among all nine tested methods, with superior AUCROC, AUPRC, and F1 scores across all negative sampling strategies (**Figure 2A-C and Figure S3**). SLxGO also demonstrated the highest ranking performance, attaining the highest NDCG@10, Precision@10, and Recall@10 values in most evaluation conditions. Performance advantages were most pronounced under expression-based and dependency-based negative sampling, where several network-dependent models, notably DDGCN and PTGNN, showed reduced performance. Across all cross-validation settings and negative sampling strategies, SLxGO obtained the highest composite classification score (0.4×C_CV1_ + 0.5×C_CV2_ + 0.1×C_CV3_) and ranking score of all evaluated models.

**Figure 2.**
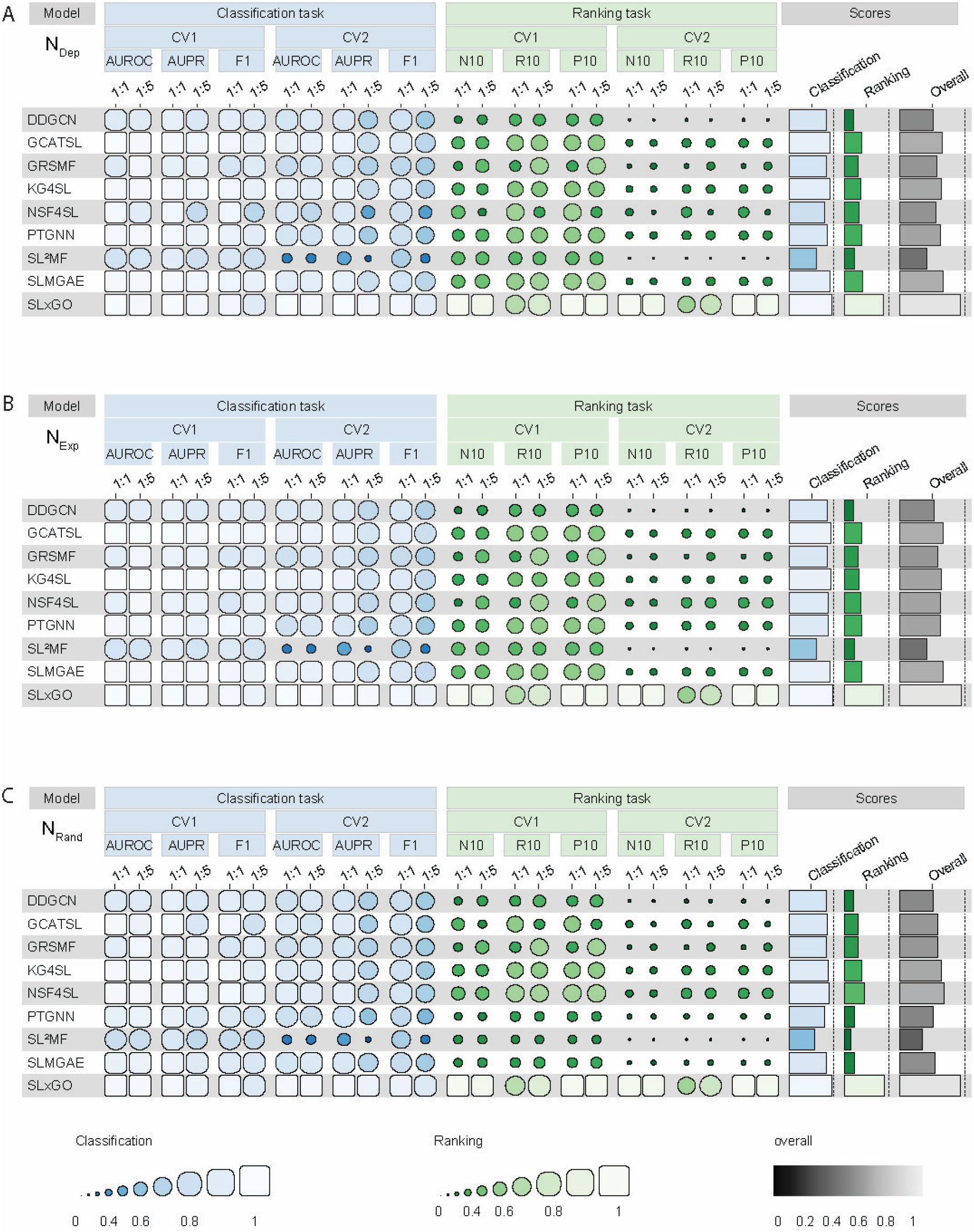
Benchmarking of SLxGO against state-of-the-art synthetic lethal prediction methods under CV1 and CV2 evaluation schemes. Comparison of SLxGO with eight existing SL prediction models (GRSMF, SL²MF, GCATSL, DDGCN, SLMGAE, NSF4SL, KG4SL, and PTGNN) following the benchmarking framework of *Feng et al.* (17) (A-C) Performance under dependency-based (NDep), expression-based (NExp), and random (NRand) negative sampling strategies. Classification performance was evaluated using AUROC, AUPRC, and F1 score, whereas ranking performance was evaluated using NDCG@10, Recall@10, and Precision@10 at both 1:1 and 5:1 positive-to-negative ratios. Summary columns show aggregate classification, ranking, and overall scores. SLxGO consistently achieves superior performance across CV1 and CV2.

The CV_3_ cross-validation setting, in which neither gene in a test pair was observed during training, represents the most stringent evaluation scenario and closely mirrors real-world applications involving newly characterized or poorly annotated genes. Under these cold-start conditions, SLxGO maintained strong classification and ranking performance across all three negative sampling strategies (**Figure 3**), whereas the majority of competing models showed substantial declines in both AUCROC and ranking metrics. Because SLxGO does not require query genes to be present in curated datasets and PPI networks, the model retains predictive capability even when interaction information is unavailable.

**Figure 3.**
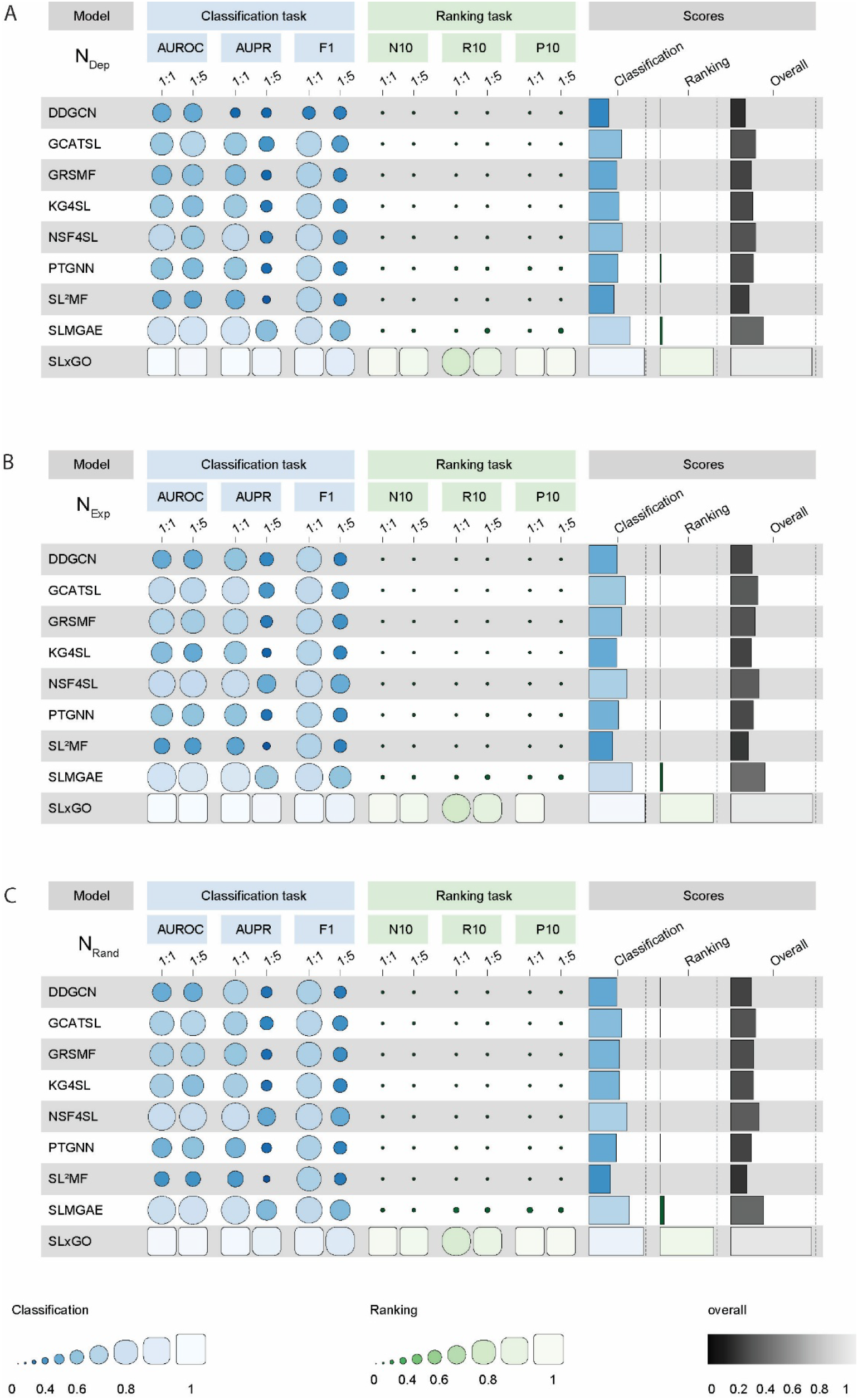
SLxGO maintains robust performance under cold-start cross-validation (CV3). (A-C) Performance under the stringent CV3 evaluation scheme, in which neither gene in a test pair was observed during training, using dependency-based (NDep), expression-based (NExp), and random (NRand) negative sampling strategies. Classification performance was evaluated using AUROC, AUPRC, and F1 score, whereas ranking performance was assessed using NDCG@10, Recall@10, and Precision@10 at both 1:1 and 5:1 positive-to-negative ratios. Summary columns show aggregate classification, ranking, and overall scores. SLxGO consistently outperforms competing methods under cold-start conditions.

To provide an overall assessment of the model performance, we further compared them by assigning equal weight to the three CV settings (**Figure S3**). Under this balanced evaluation, SLxGO achieved the highest overall classification and ranking performance, across all negative sampling conditions, demonstrating robust and consistent predictive capability across both conventional and stringent cold-start settings.

### Cell line-specific prediction reveals context-dependent SL interactions

SL interactions are highly context dependent, varying across cell types and genetic backgrounds (6, 7). To capture this context specificity, we developed SLxGO+, an extension of SLxGO that incorporates cellular context through an additional 16-dimensional gene expression embedding. This embedding was generated by applying UMAP dimensionality reduction to transcriptomic profiles from 1,206 human cell lines obtained from the Human Protein Atlas (26) **(Figure 4)**. The optimal embedding dimensionality was determined using a composite metric integrating trustworthiness, k-nearest neighbour preservation, distance correlation, and variance ratio (see Materials and Methods). Cell line-specific models were trained for six cell lines (HEK293, A375, A549, HeLa, Jurkat, and K562), each with at least 100 experimentally annotated SL and non-SL gene pairs (Table 1). Incorporating cell-specific expression information enabled SLxGO+ to achieve consistently high predictive accuracy across six cell lines, with a cumulative accuracy of 0.9584 and precision of 0.9605, indicating a low false-positive rate. Recall was more variable across datasets (cumulative recall = 0.5276), resulting in an overall F1 score of 0.6811. Performance was strongest in A375, where near-perfect accuracy (0.9980), precision (1.0000), recall (0.9954), and F1 score (0.9977) were achieved. Ranking performance was moderate overall, with a cumulative NDCG@10 of 0.5157, while top-ranked predictions showed reasonable performance, with a cumulative Precision@10 and Recall@10 values of 0.6290 and 0.5845, respectively. These results (**Table 4**) indicate that SLxGO+ prioritises highly reliable predictions while maintaining competitive ranking performance across diverse cellular contexts.

**Figure 4.**
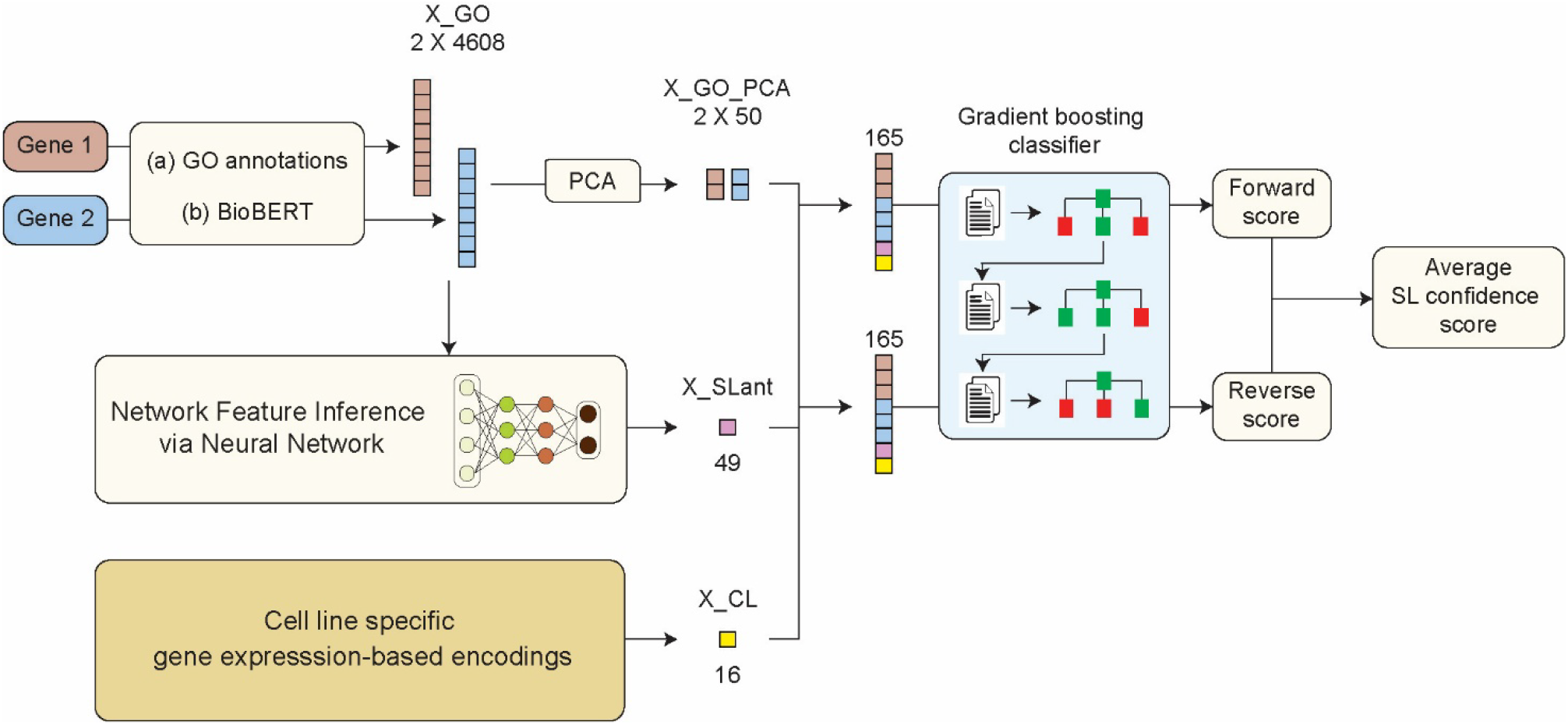
SLxGO+: a context-specific framework for synthetic lethality prediction. Schematic overview of the SLxGO+ workflow. Gene pairs are represented using Gene Ontology (GO) annotations, which are embedded using BioBERT to generate semantic feature vectors (X_GO). Principal Component Analysis (PCA) is applied to obtain a reduced representation (X_GO_PCA), while the original GO embeddings are used to infer network-like features (X_SLant) through a neural network. To enable cell line-specific prediction, a 16-dimensional gene expression-based embedding (X_CL), derived from transcriptomic profiles of 1,206 human cell lines, is incorporated. The inferred network features, reduced GO features and cell line-specific embeddings are concatenated into a 165-dimensional feature vector and used by a gradient boosting classifier to predict synthetic lethal interactions. Predictions are performed in both forward and reverse gene orientations, and the average of the two scores is reported as the final synthetic lethality confidence score.

**Table 4.**
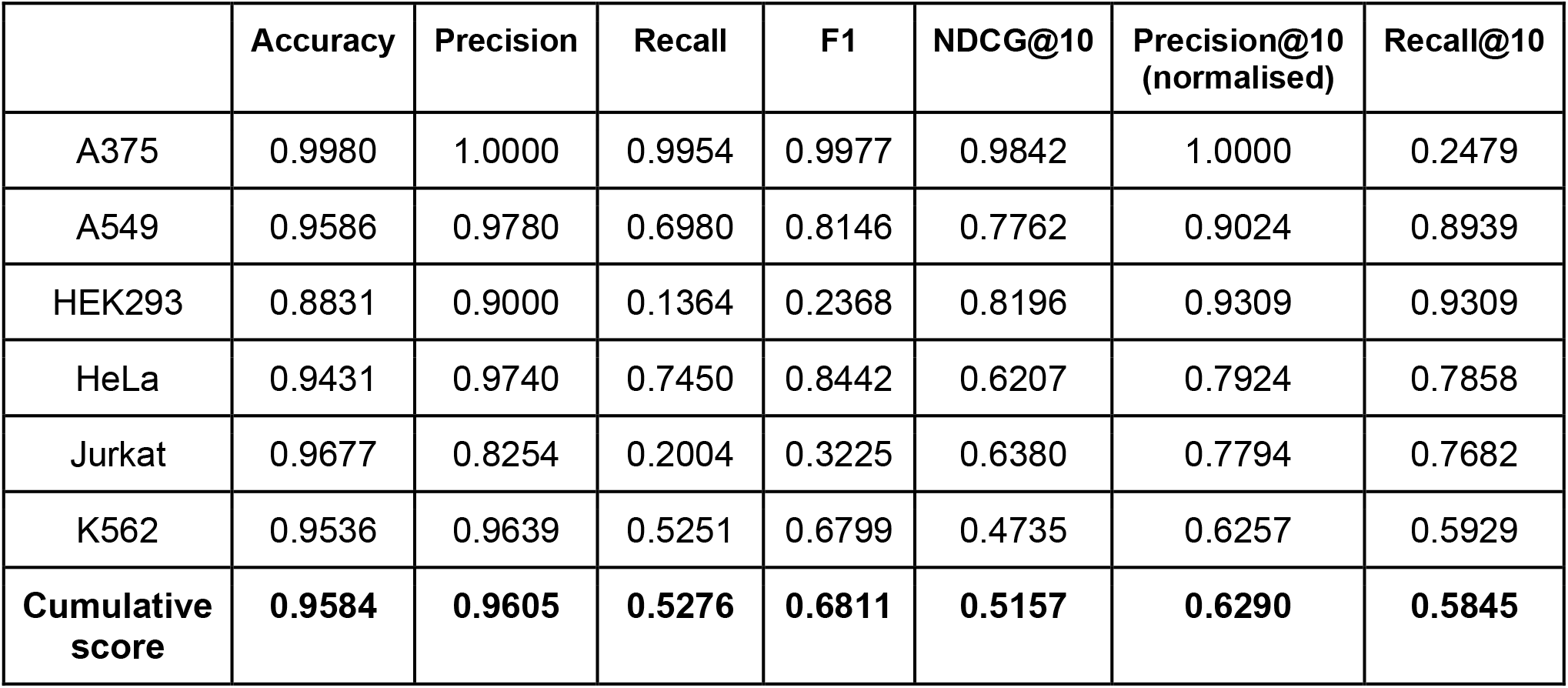
Performance of SLxGO+ on the cumulative dataset presented in Table 3.

|  | Accuracy | Precision | Recall | F1 | NDCG@10 | Precision@10<br>(normalised) | Recall@10 |
| --- | --- | --- | --- | --- | --- | --- | --- |
| A375 | 0.9980 | 1.0000 | 0.9954 | 0.9977 | 0.9842 | 1.0000 | 0.2479 |
| A549 | 0.9586 | 0.9780 | 0.6980 | 0.8146 | 0.7762 | 0.9024 | 0.8939 |
| HEK293 | 0.8831 | 0.9000 | 0.1364 | 0.2368 | 0.8196 | 0.9309 | 0.9309 |
| HeLa | 0.9431 | 0.9740 | 0.7450 | 0.8442 | 0.6207 | 0.7924 | 0.7858 |
| Jurkat | 0.9677 | 0.8254 | 0.2004 | 0.3225 | 0.6380 | 0.7794 | 0.7682 |
| K562 | 0.9536 | 0.9639 | 0.5251 | 0.6799 | 0.4735 | 0.6257 | 0.5929 |
| **Cumulative**<br>**score** | **0.9584** | **0.9605** | **0.5276** | **0.6811** | **0.5157** | **0.6290** | **0.5845** |

To systematically investigate context-dependent SL interactions, SLxGO+ was trained using the CV_1_ evaluation strategy in which predictions are generated only for gene pairs formed from combinations of genes present in the training set. Because both genes have been observed during training, this strategy provides greater confidence in predicted interactions.

To identify cell line-specific SL relationships, we selected gene pairs predicted as SL-positive in three cell lines but SL-negative in the remaining three. To further increase prediction confidence, only gene pairs with a general SLxGO prediction score greater than 0.8 were retained. This yielded 47,769 high-confidence gene pairs involving 6,247 unique genes from an initial set of 163,227 context-dependent predictions.

To prioritize therapeutically actionable interactions, we next excluded gene pairs containing essential genes. Gene essentiality was determined using Chronos scores from the DepMap database, where genes with Chronos scores greater than −0.1 across all six cell lines were classified as non-essential. This filtering step reduced the dataset to 29,702 candidate SL interactions involving 4,532 non-essential genes.

Finally, we applied a druggability filter using inhibitor annotations from Drug-Gene Interaction Database (DGIdb) (31), retaining genes with at least ten reported drug interactions. This produced a final high-confidence set of 2,641 potentially targetable SL pairs involving 529 druggable genes. The overlap and distribution of these context-specific SL interactions across the six cell lines are summarized in an UpSet plot (**Figure S4**).

### Experimental validation confirms *EFNA1*-*SLC29A1* synthetic lethality in HeLa but not in A549 cells

One of the predicted SL gene pairs identified through the systematic screening is the *EFNA1*-*SLC29A1* combination. *EFNA1* encodes for the Ephrin A1 protein involved in Ephrin signalling and *SLC29A1* encoding for the equilibrative nucleoside transporter *ENT1*. To validate the predicted *EFNA1*-*SLC29A1* SL interaction, we performed siRNA-mediated knockdown of *EFNA1* combined with pharmacological inhibition of *ENT1* using NBMPR in HeLa and A549 cells. Efficient *EFNA1* knockdown was confirmed by flow cytometry, which showed a marked reduction in *EFNA1*-associated fluorescence signal relative to scrambled siRNA controls (**Figure 5A**). Functional assessment by crystal violet staining revealed that neither *EFNA1* knockdown alone nor NBMPR treatment alone produced detectable loss of cell viability in either cell line. In contrast, combined *EFNA1* knockdown and *ENT1* inhibition produced a pronounced, dose-dependent reduction in HeLa cell viability (**Figure 5B**), consistent with a synthetic lethal interaction.

**Figure 5.**
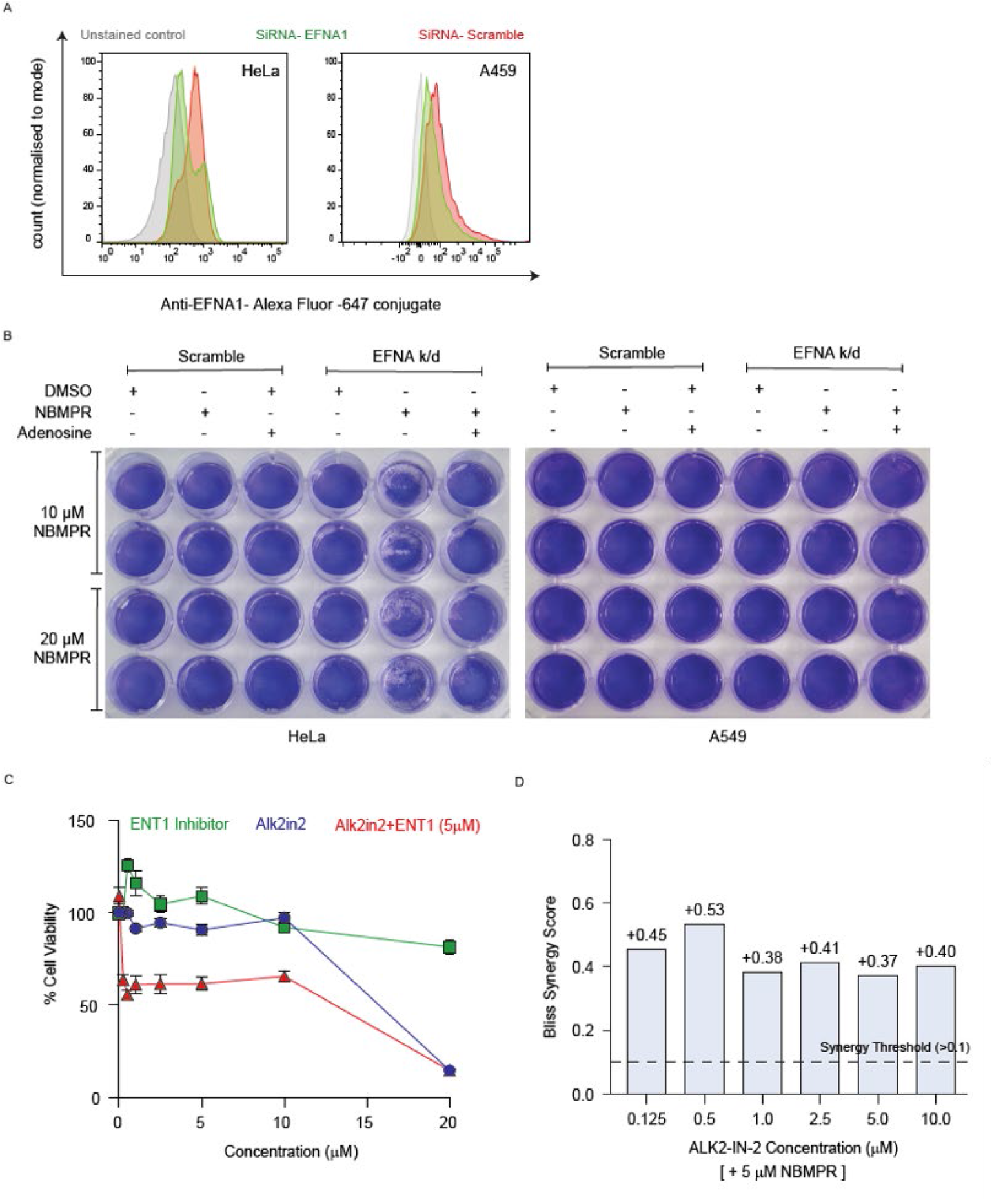
Experimental validation of the context-specific *EFNA1*–*SLC29A1* SL interaction. **(A)** Flow cytometry analysis confirming efficient *EFNA1* knockdown in HeLa cells following siRNA treatment relative to scrambled siRNA controls. **(B)** Crystal violet viability assay showing the effects of *EFNA1* knockdown and pharmacological *ENT1* inhibition using NBMPR in HeLa and A549 cells. Combined *EFNA1* knockdown and *ENT1* inhibition produced a marked, dose-dependent reduction in HeLa cell viability, whereas either perturbation alone had minimal effect. Adenosine supplementation rescued the viability defect induced by combined *EFNA1* knockdown and *ENT1* inhibition in HeLa cells, supporting a role for impaired nucleoside salvage in the observed synthetic lethal phenotype. In contrast, A549 cells showed no detectable loss of viability under identical treatment conditions, consistent with a context-specific synthetic lethal interaction between *EFNA1* and *SLC29A1*. **(C)** Combinatorial pharmacological validation of the predicted *ACVR1*-*SLC29A1* interaction. Cells were treated with an escalating dose gradient of the ACVR1 inhibitor ALK2-IN-2 (0.125 μM to 20 μM) alone or in combination with a constant 5 μM dose of the *ENT1* inhibitor NBMPR. The fixed-dose combination resulted in a synergistic reduction in cell viability compared to single-agent baselines. **(D)** Bliss Independence synergy scores calculated for the combinatorial treatments. Scores across the 0.125 μM to 10 μM ALK2-IN-2 gradient ranged from +0.29 to +0.44 (Scores > 0.1 indicate strong pharmacological synergy), highlighting the actionable nature of the SLxGO-predicted vulnerability.

To investigate the molecular basis of the predicted context-specific SL between *EFNA1* and *SLC29A1*, pathways associated with the two genes were identified based on annotations in DAVID (32) and differences in expression of genes associated with these pathways across cell lines from Human Protein Atlas (27) were analyzed. SLxGO+ predicted SL between *EFNA1* and *SLC29A1* in HeLa and HEK293 cells but not in A549 or Jurkat cells. Z-score-normalized gene expression data were used to quantify ephrin signalling, nucleoside salvage and *de novo* nucleotide synthesis in all concerned cell lines (see Methods for detailed description) (**Figure S5A**). HeLa and HEK293 cells displayed higher relative activity of nucleoside salvage and ephrin signaling pathways compared with A549 and Jurkat cells, which showed elevated de novo nucleotide synthesis activity.

To quantify these differences, a relative pathway dependency score denoting the difference between nucleoside salvage and de novo synthesis pathway activities (Salvage minus *de novo*) was defined. This metric yielded positive values in HeLa and HEK293, indicating predominant reliance on salvage-mediated nucleotide acquisition in these cells, while negative values in A549 and Jurkat indicated a dominance of *de novo* nucleotide synthesis pathway in these cells (**Figure S5B**). Together, these findings suggest that *EFNA1* perturbation increases nucleotide demand selectively in salvage-dependent cell lines, thereby creating a dependency on *ENT1*-mediated nucleoside uptake that is absent in cell lines with high baseline de novo nucleotide synthesis capacity (**Figure S5C**).

To confirm that the observed phenotype was attributable to impaired nucleoside availability, rescue experiments were performed by supplementing cultures with exogenous adenosine. Adenosine pretreatment fully rescued the viability defect in HeLa cells (**Figure 5B**), demonstrating that the synthetic lethal phenotype results from disruption of *ENT1*-mediated nucleoside transport. In contrast, A549 cells showed no detectable reduction in viability under identical treatment conditions (**Figure 5B**), confirming the cell line specificity of the *EFNA1*-*SLC29A1* interaction as predicted by SLxGO+.

### Pharmacological Validation of the *ACVR1*-*SLC29A1* Vulnerability Hub

Having established the genetic accuracy of SLxGO through the *EFNA1*-*SLC29A1* pairing, we next investigated whether our model’s predictions could be translated directly into actionable, targeted small-molecule combinations. SLxGO identified a high-confidence interaction between *ACVR1* (*ALK2*) and *SLC29A1* (*ENT1*) in HeLa cells. To validate this, we performed a combinatorial drug sensitization assay using the ACVR1 inhibitor, ALK2-IN-2 and the ENT1 inhibitor NBMPR.

Both NBMPR and ALK2-IN-2 exhibited dose-dependent reductions in cell viability when administered as single agents. To evaluate synergistic interaction, cells were exposed to an escalating gradient of ALK2-IN-2 (0.125 µM to 20 µM) in the presence of a constant, sub-lethal 5 µM dose of NBMPR (**Figure 5C**). The combinatorial treatment induced a substantially greater reduction in cell viability across the entire dose gradient compared to single-agent treatments.

To mathematically confirm this observation, we calculated synergy using the Bliss Independence model (29). The combination of ALK2-IN-2 and 5 µM NBMPR yielded highly significant Bliss synergy scores ranging from +0.37 to +0.53 at non-toxic single-agent concentrations (0.125 µM to 10 µM) (**Figure 5D**). Because a Bliss index exceeding a threshold of +0.10 denotes strict pharmacological synergy rather than mere additive cytotoxicity, these data conclusively demonstrate that pharmacological inhibition of *ENT1* highly sensitizes cells to *ACVR1* inhibition. By bridging the genetic proof-of-concept of *EFNA1* with the pharmacological exploitability of *ACVR1*, these results establish *ENT1* as a broad therapeutic vulnerability hub successfully identified by SLxGO.

### SLiGO: an open-access database for genome-scale SL predictions

To facilitate community access to SLxGO and SLxGO+ predictions, we developed SLiGO (Gene Ontology-driven Synthetic Lethal Interactions; https://databases.iisertvm.ac.in/sligo/), a freely accessible web-based database covering SL predictions for 30 million human gene pairs across six cell lines (**Figure 6A-B**). The SLiGO platform provides gene-centric and gene pair query interfaces returning predicted SL partners together with prediction confidence scores, GO-derived annotations, and links to external resources (GeneCards, UniProt). Results can be browsed interactively or downloaded in bulk. SLiGO is an open-access resource that does not require any registration and will be updated periodically as model refinements become available.

**Figure 6.**
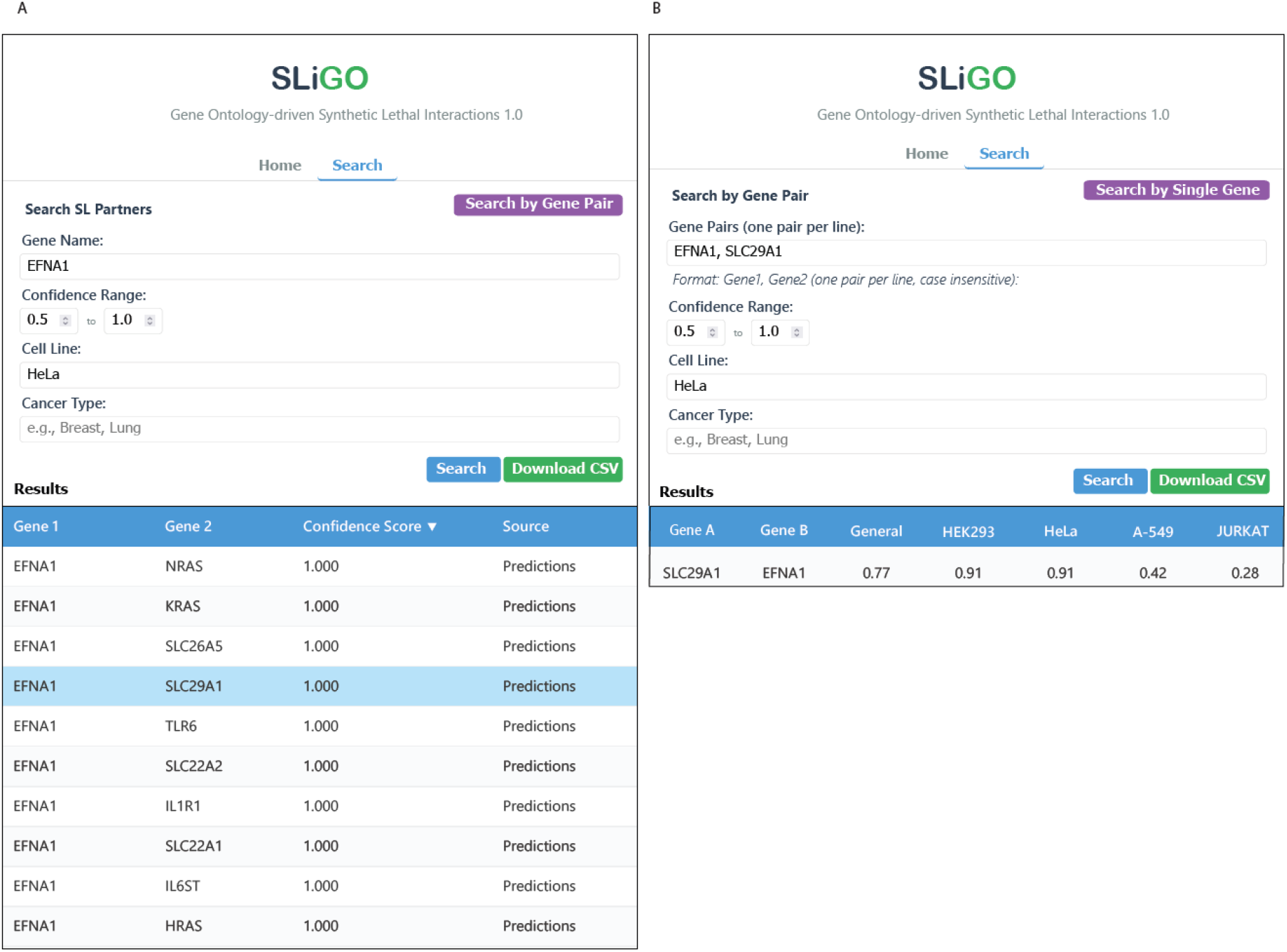
SLiGO: an open-access database for genome-scale synthetic lethality predictions. **(A)** Gene-centric search interface showing predicted SL partners of *EFNA1* in HeLa cells. **(B)** Pairwise query interface displaying cell-line-specific confidence scores for the *EFNA1*-*SLC29A1* gene pair across four cell lines.

## Discussion

In this study, we introduce SLxGO, a gradient boosting-based model that predicts synthetic lethality directly from GO-derived semantic representations. To establish the validity of this approach, we conducted a rigorous comparative benchmarking evaluation against eight state-of-the-art, network-dependent methods. SLxGO demonstrated consistent performance gains across all baselines, particularly under stringent cross-validation paradigms (CV_3_) designed to simulate ‘cold-start’ conditions involving completely unseen genes. These benchmarking results indicate that much of the predictive information previously attributed to explicit network topology is already latent within the functional annotation space. While the use of a tree-based classifier architecture may partially contribute to improvements in ranking metrics (33), the model’s sustained superiority under strict CV3 conditions confirms that the benchmarked performance is driven by the robust, transferable predictive information encoded within the GO semantic embeddings themselves, rather than data leakage or algorithmic bias. Furthermore, our cell line-specific extension demonstrates that incorporating transcriptional state meaningfully reshapes predicted SL landscapes, consistent with growing evidence that SL is determined as much by cellular physiology as by genetic architecture (6, 7). Gene pairs ranked highly in one cellular context are frequently deprioritized in another, underscoring the inadequacy of context-agnostic SL prediction for translational applications where the relevant cellular background is a critical clinical variable.

A pivotal insight arising from our benchmarking and subsequent SHAP feature attribution analysis is that inferred network-like features contribute only modestly to model performance compared to GO semantic features. This disparity demonstrates that GO-derived semantic representations inherently encapsulate the higher-order functional relationships required for robust, genome-scale SL inference, mitigating the need for explicit topological frameworks. This observation extends far beyond SL modeling; it implies that annotation-based semantic representations can effectively decouple predictive performance from explicit network architectures across diverse computational biology tasks. Consequently, this approach offers a viable path forward for resolving the ‘cold-start’ problem in gene function prediction, allowing accurate modeling of genes that currently lack interaction data within biological knowledge graphs.

The experimental validation of the *EFNA1*-*SLC29A1* interaction provides strong empirical evidence that the high performance metrics observed during benchmarking translate directly to true biological dependencies. Combined perturbation of *EFNA1* and *SLC29A1* selectively induces lethality in HeLa but not A549 cells, consistent with model predictions. This context-specificity is mechanistically coherent with lineage-specific differences in nucleotide metabolism. HeLa cells exhibit higher dependence on the nucleoside salvage pathway, in which *SLC29A1* plays a central role, while A549 cells appear to compensate through elevated *de novo* nucleotide synthesis (34). *EFNA1* perturbation likely amplifies dependence on *SLC29A1*-mediated salvage selectively in cells lacking sufficient compensatory capacity. Importantly, this mechanistic logic is not explicitly encoded in SLxGO, which has no access to pathway structure or metabolic flux data. Notably, these functional relationships captured from GO semantic embeddings manifested as experimentally validated context-specific vulnerabilities.

Beyond genetic validation, the ultimate utility of *in silico* synthetic lethal (SL) predictions lies in their translational potential. Our pharmacological validation of the *ACVR1*-*SLC29A1* interaction demonstrates that SLxGO can identify therapeutically actionable, cross-pathway vulnerabilities. The profound synergy observed upon dual inhibition of *ACVR1* and *ENT1* highlights a critical metabolic bottleneck: oncogenic signaling cascades frequently exacerbate replication stress and deplete intracellular nucleotide pools, thereby rendering cells highly dependent on the *ENT1*-mediated nucleoside salvage pathway. By accurately predicting such intricate metabolic dependencies independent of protein-protein interaction (PPI) networks, SLxGO overcomes the inherent data biases and topological constraints of existing models identified in our benchmarking. Importantly, the successful transition from a GO-based computational prediction to a highly synergistic pharmacological intervention underscores SLxGO as a robust, scalable framework for rational combinatorial drug discovery in precision oncology.

Several limitations of the current framework warrant consideration. SLxGO predictions are bounded by the completeness of GO annotations, which remain sparse for poorly characterized genes and may lag behind emerging biological knowledge (18). Although this is precisely the gene space where network-based methods also fail, it represents an unresolved challenge for annotation-driven approaches broadly. A more fundamental constraint is that comprehensive GO annotation coverage currently exists primarily for human genes (35–37), limiting the direct applicability of SLxGO to other species without substantial retraining on species-specific annotation data. This restricts the framework to human cancer contexts and precludes straightforward extension to model organisms such as yeast or mouse, where SL screens have generated valuable experimental data (38, 39). Additionally, while the neural network effectively learns transferable network-derived topological representations from functional semantics, the biological interpretability of these inferred features warrants further investigation.

To facilitate community use, all SLxGO predictions are freely accessible through SLiGO (https://databases.iisertvm.ac.in/sligo/), an open-access database covering 30 million human gene pairs across six cell lines. Future developments will focus on extending the predictions to CV_2_ and CV_3_ conditions significantly expanding the database to over 200 million human gene pairs. In addition, incorporating a gene regulatory network (GRN) structure-based layer that captures transcriptional regulatory relationships across diverse cellular and species contexts, providing a complementary source of biological signal to further extend SLxGO predictions beyond the current dependency on human GO annotation coverage. This would enable broader applicability across additional cell lines and model organisms where SL screens have generated experimental ground truth data. Additional planned developments include integration of pathway-level annotations and single-cell expression profiles to refine context-specific predictions and expand coverage of understudied genes. In conclusion, SLxGO and SLiGO together provide a scalable, network-independent resource for genome-scale, context-specific disease vulnerability mapping that substantially expands the searchable SL landscape beyond the limits of existing interaction maps.

## Data availability

All data supporting the findings of this study are available within the Article, Supplementary Information, Source Data files, and the project’s GitHub repository. Additional materials are available from the corresponding author upon reasonable request.

## Code availability

Custom scripts used for data preprocessing, feature generation, model training, benchmarking, and downstream computational analyses are available at the GitHub repository (https://github.com/sshameer/SLxGO2026).

## Use of Artificial Intelligence Tools

ChatGPT (OpenAI) and Gemini (Google LLC) were used during manuscript preparation to assist with language editing and improve readability. All scientific content, analyses, interpretations, and conclusions were verified by the authors, who take full responsibility for the final manuscript.

## Acknowledgements

This work was supported by the DBT-Wellcome Trust India Alliance Intermediate Fellowship (IA/I/23/2/506998) to K.V., Anusandhan National Research Foundation (SERB-ANRF) (ARG/2025/011068/LS) to S.S., and intramural funding from the Indian Institute of Science Education and Research Thiruvananthapuram (IISER Thiruvananthapuram) to K.V, S.S and S.B. P.B and P.S received IISER Thiruvananthapuram institute PhD fellowship throughout the study.

## Author Contributions

PB, AL and PS performed experiments. PB, PS, SS, SB and KV analyzed data. SS, SB and KV supervised the research. PB, SS and KV wrote the paper with input from all other authors.

## References

1. Turk, A.A. and Wisinski, K.B. (2018) PARP inhibitors in breast cancer: Bringing synthetic lethality to the bedside. Cancer, 124, 2498–2506.

2. Setton, J., Zinda, M., Riaz, N., Durocher, D., Zimmermann, M., Koehler, M., Reis-Filho, J.S. and Powell, S.N. (2021) Synthetic lethality in cancer therapeutics: The next generation. Cancer Discov., 11, 1626–1635.

3. Schäffer, A.A., Chung, Y., Kammula, A.V., Ruppin, E. and Lee, J.S. (2024) A systematic analysis of the landscape of synthetic lethality-driven precision oncology. Med (N. Y.), 5, 73–89.e9.

4. Gonçalves, E., Ryan, C.J. and Adams, D.J. (2026) Synthetic lethality in cancer drug discovery: challenges and opportunities. Nat Rev Drug Discov, 25, 22–38.

5. Tang, S., Gökbağ, B., Fan, K., Shao, S., Huo, Y., Wu, X., Cheng, L. and Li, L. (2022) Synthetic lethal gene pairs: Experimental approaches and predictive models. Front. Genet., 13, 961611.

6. Ku, A.A., Hu, H.-M., Zhao, X., Shah, K.N., Kongara, S., Wu, D., McCormick, F., Balmain, A. and Bandyopadhyay, S. (2020) Integration of multiple biological contexts reveals principles of synthetic lethality that affect reproducibility. Nat Commun, 11, 2375.

7. Chang, L., Shaw, K., Vazquez, F. and Sellers, W.R. (2026) Context-dependent synthetic lethality - an emerging precision therapeutic approach. Nat Rev Cancer, 10.1038/s41568-026-00929-9.

8. Jerby-Arnon, L., Pfetzer, N., Waldman, Y.Y., McGarry, L., James, D., Shanks, E., Seashore-Ludlow, B., Weinstock, A., Geiger, T., Clemons, P.A., et al. (2014) Predicting cancer-specific vulnerability via data-driven detection of synthetic lethality. Cell, 158, 1199–1209.

9. Benstead-Hume, G., Chen, X., Hopkins, S.R., Lane, K.A., Downs, J.A. and Pearl, F.M.G. (2019) Predicting synthetic lethal interactions using conserved patterns in protein interaction networks. PLoS Comput. Biol., 15, e1006888.

10. Huang, J., Wu, M., Lu, F., Ou-Yang, L. and Zhu, Z. (2019) Predicting synthetic lethal interactions in human cancers using graph regularized self-representative matrix factorization. BMC Bioinformatics, 20, 657.

11. Liu, Y., Wu, M., Liu, C., Li, X.-L. and Zheng, J. (2020) SL2MF: Predicting synthetic lethality in human cancers via logistic matrix factorization. IEEE/ACM Trans. Comput. Biol. Bioinform., 17, 748–757.

12. Long, Y., Wu, M., Liu, Y., Zheng, J., Kwoh, C.K., Luo, J. and Li, X. (2021) Graph contextualized attention network for predicting synthetic lethality in human cancers. Bioinformatics, 37, 2432– 2440.

13. Cai, R., Chen, X., Fang, Y., Wu, M. and Hao, Y. (2020) Dual-dropout graph convolutional network for predicting synthetic lethality in human cancers. Bioinformatics, 36, 4458–4465.

14. Kosoglu, K., Aydin, Z., Tuncbag, N., Gursoy, A. and Keskin, O. (2023) Structural coverage of the human interactome. Brief Bioinform, 25.

15. Seale, C., Tepeli, Y. and Gonçalves, J.P. (2022) Overcoming selection bias in synthetic lethality prediction. Bioinformatics, 38, 4360–4368.

16. Lee, J., Yoon, W., Kim, S., Kim, D., Kim, S., So, C.H. and Kang, J. (2020) BioBERT: a pre-trained biomedical language representation model for biomedical text mining. Bioinformatics, 36, 1234–1240.

17. Feng, Y., Long, Y., Wang, H., Ouyang, Y., Li, Q., Wu, M. and Zheng, J. (2024) Benchmarking machine learning methods for synthetic lethality prediction in cancer. Nat. Commun., 15, 9058.

18. Gene Ontology Consortium, Aleksander, S.A., Balhoff, J., Carbon, S., Cherry, J.M., Drabkin, H.J., Ebert, D., Feuermann, M., Gaudet, P., Harris, N.L., et al. (2023) The Gene Ontology knowledgebase in 2023. Genetics, 224.

19. Ashburner, M., Ball, C.A., Blake, J.A., Botstein, D., Butler, H., Cherry, J.M., Davis, A.P., Dolinski, K., Dwight, S.S., Eppig, J.T., et al. (2000) Gene ontology: tool for the unification of biology. The Gene Ontology Consortium. Nat. Genet., 25, 25–29.

20. Arafeh, R., Shibue, T., Dempster, J.M., Hahn, W.C. and Vazquez, F. (2025) The present and future of the Cancer Dependency Map. Nat Rev Cancer, 25, 59–73.

21. Chen, X., Cai, R., Huang, Z., Li, Z., Zheng, J. and Wu, M. (2025) Interpretable high-order knowledge graph neural network for predicting synthetic lethality in human cancers. Brief. Bioinform., 26.

22. Wang, S., Feng, Y., Liu, X., Liu, Y., Wu, M. and Zheng, J. (2022) NSF4SL: negative-sample-free contrastive learning for ranking synthetic lethal partner genes in human cancers. Bioinformatics, 38, ii13–ii19.

23. Wang, S., Xu, F., Li, Y., Wang, J., Zhang, K., Liu, Y., Wu, M. and Zheng, J. (2021) KG4SL: knowledge graph neural network for synthetic lethality prediction in human cancers. Bioinformatics, 37, i418–i425.

24. Long, Y., Wu, M., Liu, Y., Fang, Y., Kwoh, C.K., Chen, J., Luo, J. and Li, X. (2022) Pre-training graph neural networks for link prediction in biomedical networks. Bioinformatics, 38, 2254– 2262.

25. Guo, J., Liu, H. and Zheng, J. (2016) SynLethDB: synthetic lethality database toward discovery of selective and sensitive anticancer drug targets. Nucleic Acids Res., 44, D1011–7.

26. Uhlén, M., Fagerberg, L., Hallström, B.M., Lindskog, C., Oksvold, P., Mardinoglu, A., Sivertsson, Å., Kampf, C., Sjöstedt, E., Asplund, A., et al. (2015) Proteomics. Tissue-based map of the human proteome. Science, 347, 1260419.

27. Jin, H., Zhang, C., Zwahlen, M., von Feilitzen, K., Karlsson, M., Shi, M., Yuan, M., Song, X., Li, X., Yang, H., et al. (2023) Systematic transcriptional analysis of human cell lines for gene expression landscape and tumor representation. Nat Commun, 14, 5417.

28. Subramanian, A., Tamayo, P., Mootha, V.K., Mukherjee, S., Ebert, B.L., Gillette, M.A., Paulovich, A., Pomeroy, S.L., Golub, T.R., Lander, E.S., et al. (2005) Gene set enrichment analysis: a knowledge-based approach for interpreting genome-wide expression profiles. Proc Natl Acad Sci U S A, 102, 15545–15550.

29. Bliss, C.I. (1939) The toxicity of poisons applied jointly1. Ann. Appl. Biol., 26, 585–615.

30. Hao, Z., Wu, D., Fang, Y., Wu, M., Cai, R. and Li, X. (2021) Prediction of synthetic lethal interactions in human cancers using Multi-view Graph Auto-Encoder. IEEE J. Biomed. Health Inform., 25, 4041–4051.

31. Cannon, M., Stevenson, J., Stahl, K., Basu, R., Coffman, A., Kiwala, S., McMichael, J.F., Kuzma, K., Morrissey, D., Cotto, K., et al. (2024) DGIdb 5.0: rebuilding the drug-gene interaction database for precision medicine and drug discovery platforms. Nucleic Acids Res, 52, D1227– D1235.

32. Sherman, B.T., Panzade, G., Dotrang, T., Hao, M., Xu, L., Li, X., Baseler, M.W., Lane, H.C., Imamichi, T. and Chang, W. (2026) DAVID: a web server for functional annotation and functional enrichment analysis of gene lists (2025 update). Nucleic Acids Res, 10.1093/nar/gkag470.

33. Qin, Z., Yan, L., Zhuang, H. and Tay, Y. (2021) Are neural rankers still outperformed by gradient boosted decision trees? International.

34. Lane, A.N. and Fan, T.W.-M. (2015) Regulation of mammalian nucleotide metabolism and biosynthesis. Nucleic Acids Res., 43, 2466–2485.

35. Huntley, R.P., Sawford, T., Martin, M.J. and O’Donovan, C. (2014) Understanding how and why the Gene Ontology and its annotations evolve: the GO within UniProt. Gigascience, 3, 4.

36. Liu, M. and Thomas, P.D. (2019) GO functional similarity clustering depends on similarity measure, clustering method, and annotation completeness. BMC Bioinformatics, 20, 155.

37. Buza, T.J., McCarthy, F.M., Wang, N., Bridges, S.M. and Burgess, S.C. (2008) Gene Ontology annotation quality analysis in model eukaryotes. Nucleic Acids Res., 36, e12.

38. O’Neil, N.J., Bailey, M.L. and Hieter, P. (2017) Synthetic lethality and cancer. Nat. Rev. Genet., 18, 613–623.

39. Costanzo, M., VanderSluis, B., Koch, E.N., Baryshnikova, A., Pons, C., Tan, G., Wang, W., Usaj, M., Hanchard, J., Lee, S.D., et al. (2016) A global genetic interaction network maps a wiring diagram of cellular function. Science, 353, aaf1420.

