## Supplementary Figures for "Genome-scale prediction of context-specific synthetic lethality beyond protein interaction networks"

Supplementary Figure 1

A

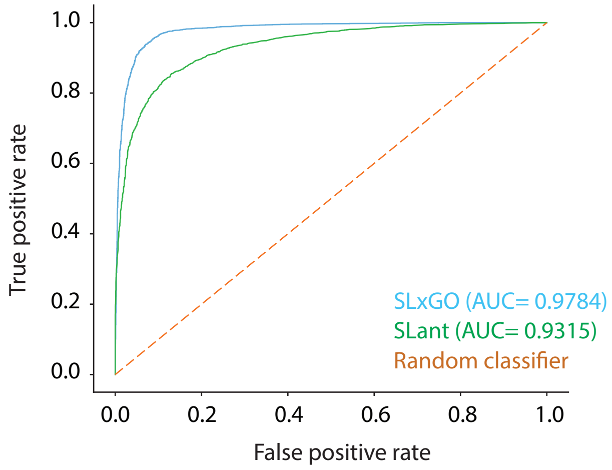

Supplementary Figure 2

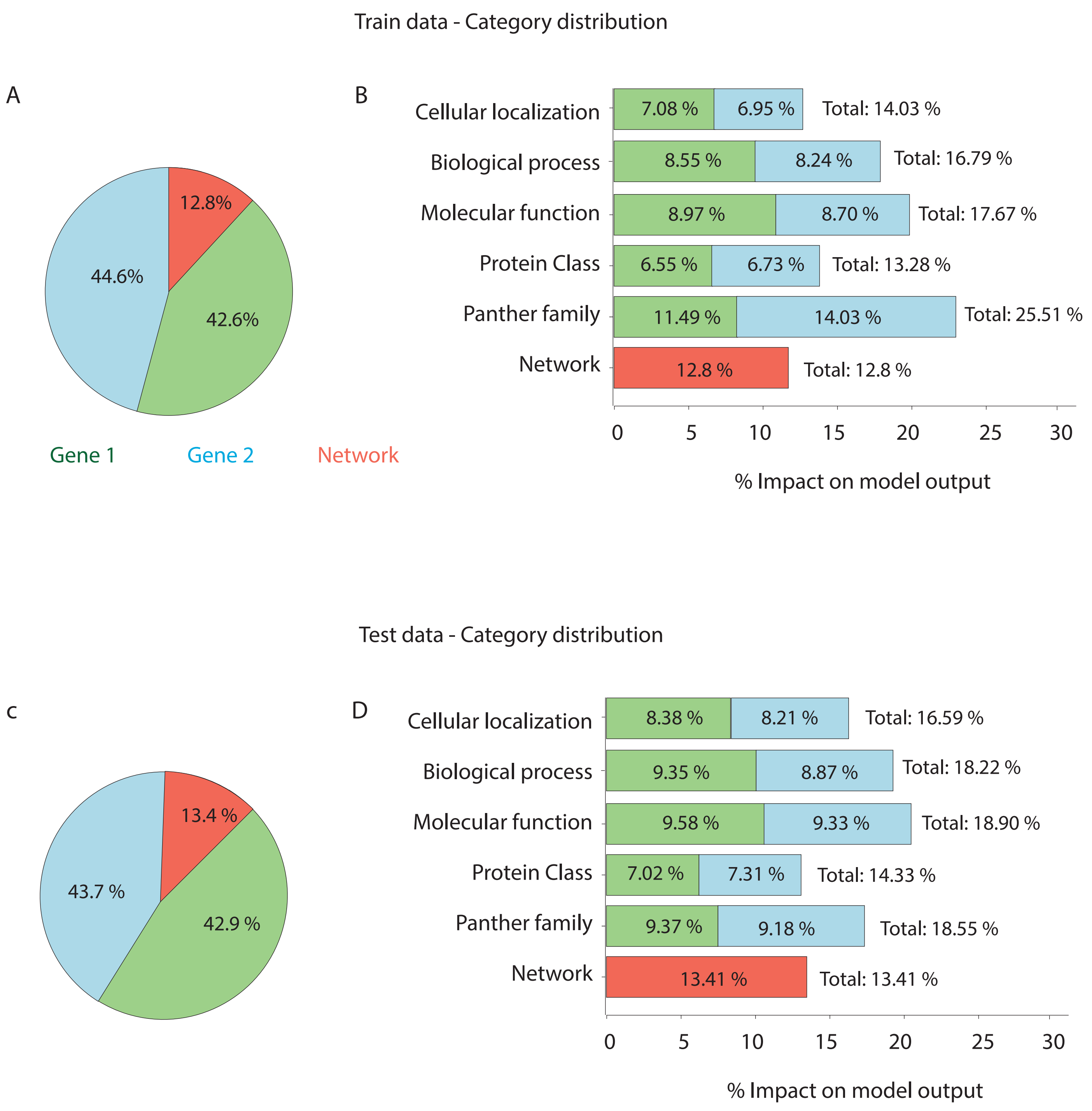

Supplementary Figure 3

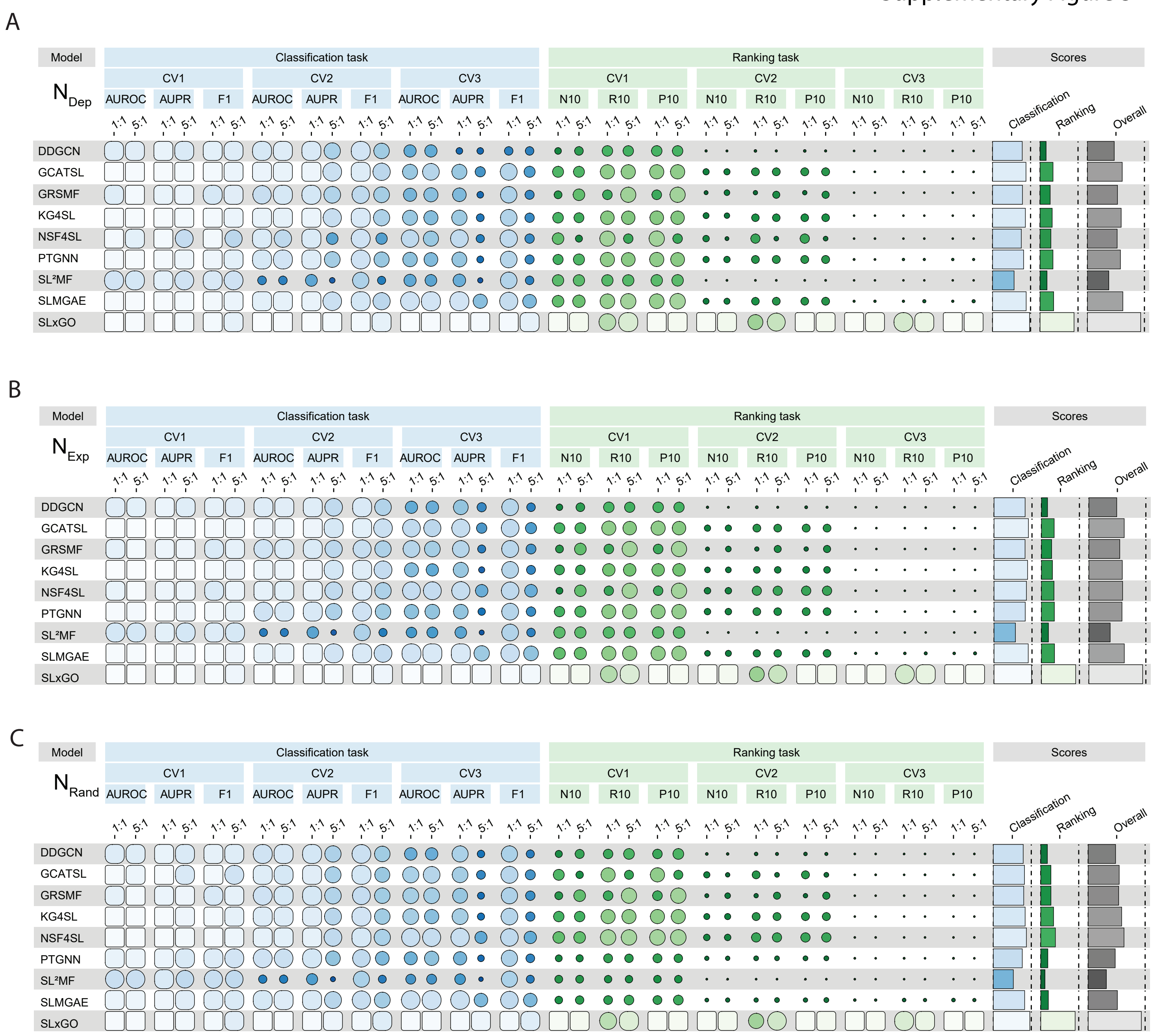

A

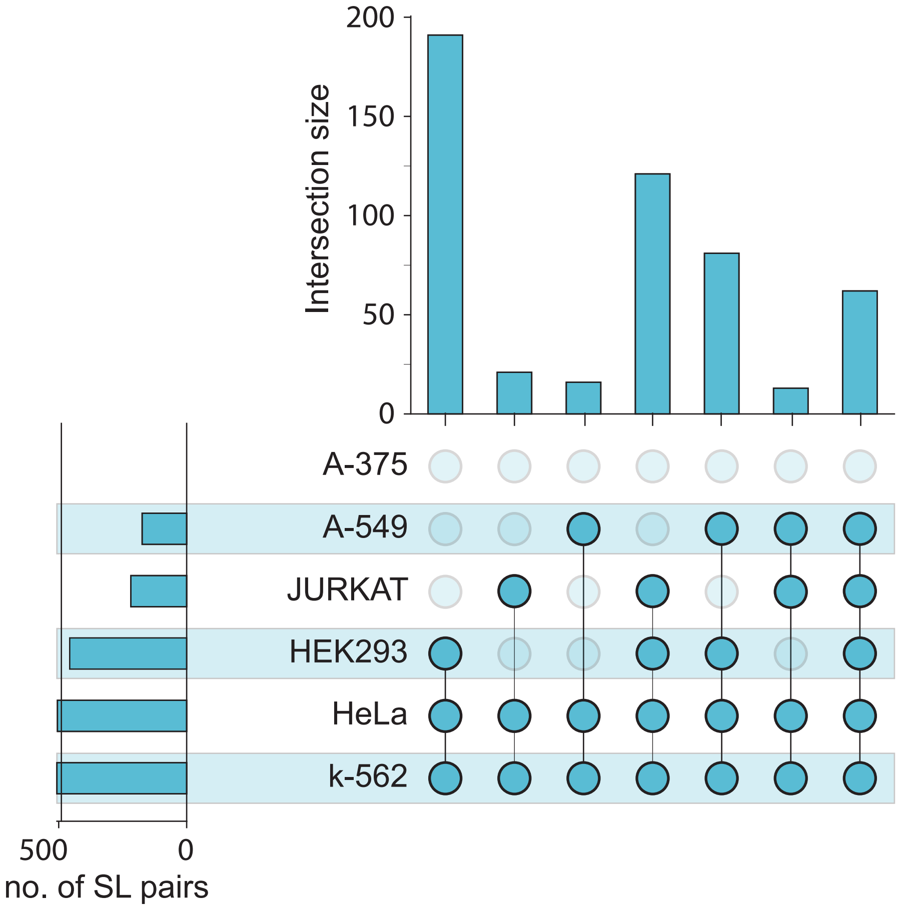
Intersection size
200
150
100
50
0
A-375
A-549
JURKAT
HEK293
HeLa
k-562
500
0
no. of SL pairs

B

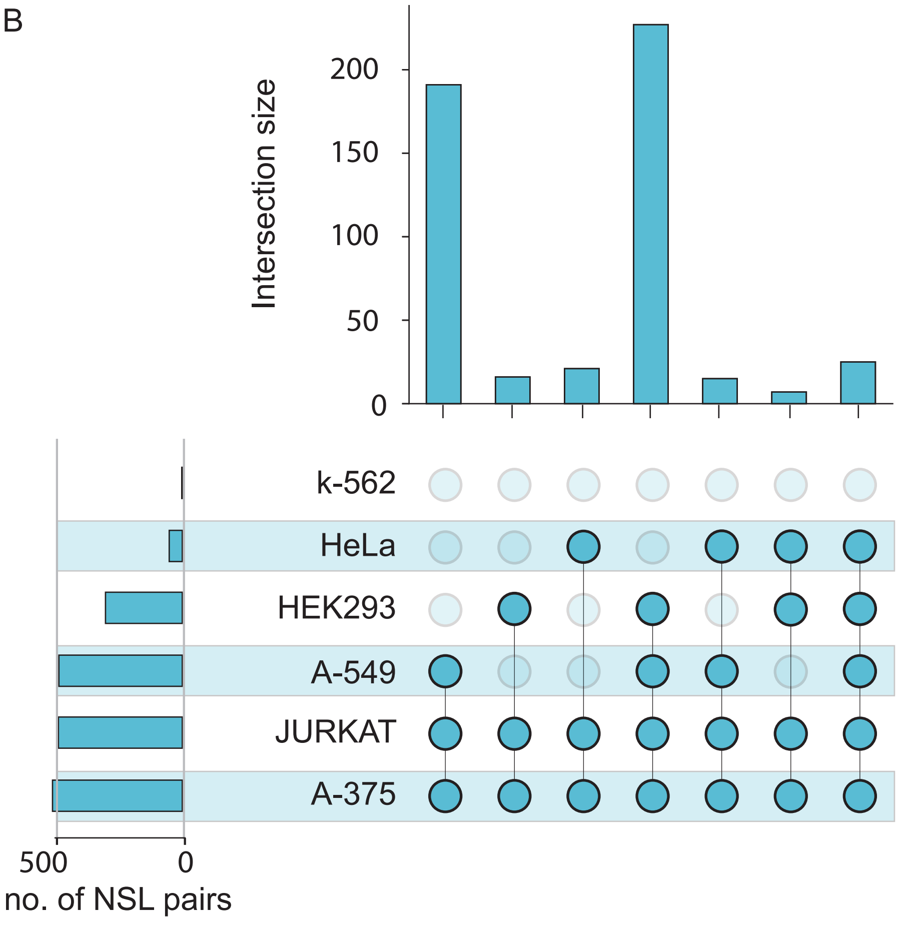
Intersection size
200
150
100
50
0
k-562
HeLa
HEK293
A-549
JURKAT
A-375
500
0
no. of NSL pairs

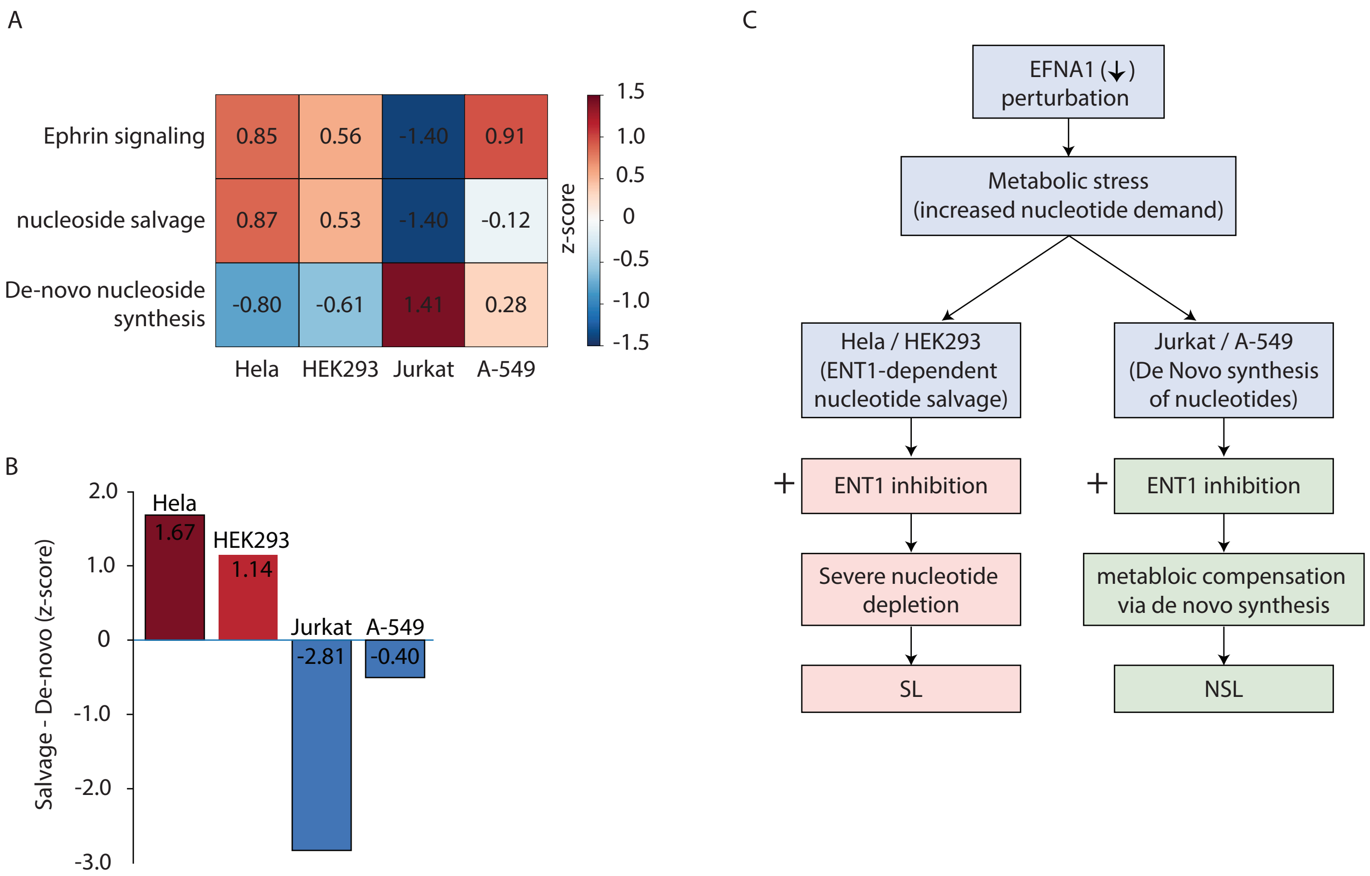
A
Ephrin signaling
nucleoside salvage
De-novo nucleoside synthesis
0.85
0.56
-1.40
0.91
0.87
0.53
-1.40
-0.12
-0.80
-0.61
1.41
0.28
Hela
HEK293
Jurkat
A-549
z-score
1.5
1.0
0.5
0
-0.5
-1.0
-1.5
B
Salvage - De-novo (z-score)
2.0
1.0
0
-1.0
-2.0
-3.0
Hela
1.67
HEK293
1.14
Jurkat
-2.81
A-549
-0.40
C
EFNA1 (↓) perturbation
Metabolic stress (increased nucleotide demand)
Hela / HEK293 (ENT1-dependent nucleotide salvage)
Jurkat / A-549 (De Novo synthesis of nucleotides)
+
ENT1 inhibition
+
ENT1 inhibition
Severe nucleotide depletion
metabloic compensation via de novo synthesis
SL
NSL

**Supplementary Figure S1. SLxGO outperforms SLant on network-covered gene pairs.** Receiver operating characteristic (ROC) curves comparing predictive performance of SLxGO and SLant on a balanced dataset of 12,690 SLant-compatible gene pairs. SLxGO showed improved discriminative performance relative to SLant, achieving a higher AUCROC.

**Supplementary Figure S2. SHAP feature contribution analysis.** SHAP analysis of feature contributions in SLxGO. **(A, B)** Feature importance distributions in the training dataset showing contributions from network-derived features, Gene1 features, and Gene2 features. **(C, D)** Corresponding feature contributions in the test dataset. GO-derived semantic features from both genes dominate model predictions, while inferred network features contribute a smaller but consistent fraction.

**Supplementary Figure S3. Benchmarking of SLxGO using equal weighting across cross-validation schemes.** Performance of SLxGO and eight benchmark methods was re-evaluated using equal weighting of the three cross-validation settings (CV1:CV2:CV3 = 1:1:1) instead of the weighting scheme (4:5:1) used in the primary analysis. Results are shown for (A) dependency-based (NDep), (B) expression-based (NExp), and (C) random (NRand) negative sampling strategies. Classification (AUCROC, AUPRC, and F1) and ranking (NDCG@10, Recall@10, and Precision@10) metrics are presented for both weighting schemes together with the corresponding aggregate classification, ranking, and overall scores.

**Supplementary Figure S4. UpSet plot of filtered SLxGO+ predictions.** UpSet plots were used as an alternative to Venn diagrams to visualize the overlap of filtered positive **(A)** and negative **(B)** SLxGO+ predictions across the six cell lines. Horizontal bars show the total number of predictions in each cell line, and vertical bars show the size of each intersection, with connected dots indicating the contributing cell line combinations.

**Supplementary Figure S5. Pathway activity analysis reveals context-specific dependency on nucleoside salvage for the EFNA1-SLC29A1 SL interaction. (A)** Heatmap of Z-score-normalized pathway activity across four cancer cell lines (HeLa, HEK293, A549, Jurkat) for three gene sets: ephrin signaling, nucleoside salvage, and de novo nucleotide synthesis. Values represent mean Z-score across genes in each pathway per cell line. **(B)** Relative pathway dependency score (Salvage − *De-novo* activity) for each cell line. Positive values indicate predominant reliance on nucleoside salvage; negative values indicate *de-novo* pathway dominance. HeLa and HEK293 show positive scores, consistent with SLxGO+-predicted SL between EFNA1 and SLC29A1; A549 and Jurkat show negative scores, consistent with predicted non-SL. **(C)** Schematic model illustrating the proposed mechanism: EFNA1 perturbation increases nucleotide demand, creating a selective dependency on ENT1-mediated nucleoside import in salvage-dependent cell lines.

**Supplementary Data S1. Benchmarking result of eight previously established SL prediction models**
